# Spatial-ATAC-seq: spatially resolved chromatin accessibility profiling of tissues at genome scale and cellular level

**DOI:** 10.1101/2021.06.06.447244

**Authors:** Yanxiang Deng, Marek Bartosovic, Sai Ma, Di Zhang, Yang Liu, Xiaoyu Qin, Graham Su, Mina L. Xu, Stephanie Halene, Joseph E. Craft, Gonçalo Castelo-Branco, Rong Fan

## Abstract

Cellular function in tissue is dependent upon the local environment, requiring new methods for spatial mapping of biomolecules and cells in the tissue context. The emergence of spatial transcriptomics has enabled genome-scale gene expression mapping, but it remains elusive to capture spatial epigenetic information of tissue at cellular level and genome scale. Here we report on spatial-ATAC-seq: spatially resolved chromatin accessibility profiling of tissue section via next-generation sequencing by combining *in situ* Tn5 transposition chemistry and microfluidic deterministic barcoding. Spatial chromatin accessibility profiling of mouse embryos delineated tissue region-specific epigenetic landscapes and identified gene regulators implicated in the central nerve system development. Mapping the accessible genome in human tonsil tissue with 20μm pixel size revealed spatially distinct organization of immune cell types and states in lymphoid follicles and extrafollicular zones. This technology takes spatial biology to a new realm by enabling spatially resolved epigenomics to improve our understanding of cell identity, state, and fate decision in relation to epigenetic underpinnings in development and disease.

## MAIN TEXT

Single cell sequencing presents a tangible way to define, in an unbiased manner, cell types and states^1–3^, but the tissue dissociation process unfortunately leads to the loss of spatial context. The field of spatial transcriptomics emerged to address this challenge and to transform how we delineate cellular functions and states in the native tissue environment^4^. It includes imaging-based approaches such as multiplexed single-molecule fluorescent *in situ* hybridization^5–8^, which evolved from detecting a handful of genes to thousands^9–11^, and Next-Generation Sequencing(NGS)-based approaches for unbiased genome-wide gene expression mapping at cellular level^12–14^. To investigate the mechanisms underlying spatial organization of different cell types and functions in the tissue context, it is highly desired to examine not only gene expression but also epigenetic underpinnings such as chromatin accessibility^15–17^ in a spatially resolved manner to uncover the causative relationship determining what drives tissue organization and function. To date, it remains elusive to spatially map epigenetic states such as chromatin accessibility directly in a tissue section at genome scale and cellular level.

Genome-wide profiling of chromatin accessibility by sequencing using a Tn5 transposition chemistry (ATAC-seq) was developed to detect all accessible genomic loci, which was further applied to single cells^15–18^. It was also demonstrated to image chromatin accessibility in fixed cells using fluorescence-labeled DNA oligomers assembled in Tn5 (ATACsee)^19^, suggesting the feasibility to profile chromatin accessibility *in situ* in a tissue section. Microdissecting tissues from specific regions using microbiopsy punching followed by single-cell ATAC-seq allowed to profile accessible chromatin of single cells from a region of interest defined by micropunching^20^. However, spatially resolved genome-scale chromatin accessibility mapping over a tissue section at cellular level has not been possible. We previously developed DBiT-seq for spatially resolved multi-omics sequencing via microfluidic barcoding of RNAs or proteins directly in tissue^14^. Herein, we applied this spatial barcoding scheme to labeling DNA oligomers that were inserted to the accessible genomic loci by Tn5 transposition followed by high throughout sequencing to realize spatial-ATAC-seq: high-spatial-resolution genome-wide mapping of chromatin accessibility in tissue at cellular level. The results from mouse embryos delineated the tissue region-specific epigenetic landscapes and gene regulators implicated in the central nerve system development. Spatial-ATAC-seq of human tonsil tissue with 20μm pixel size revealed spatially distinct organization of immune cell types and states in relation to lymphoid follicles and extrafollicular zones. This technology adds a new dimension to the study of spatial biology by bringing spatial epigenomics to the field and may find a wide range of applications in normal development and pathogenesis studies.

### Spatial chromatin accessibility sequencing design and workflow

We present spatial-ATAC-seq for mapping chromatin accessibility in a tissue section at cellular level via combining the strategy of microfluidic deterministic barcoding in tissue^21^ and the chemistry of the assay for transposase-accessible chromatin^15,22^ (Fig. 1a and Fig. S1). The main workflow for spatial ATAC-seq is shown in Fig. 1a. The fresh frozen tissue section on a standard aminated glass slide was fixed with formaldehyde. Tn5 transposition was then performed and the adapters containing a ligation linker were inserted to transposase accessible genomic DNA loci. Afterwards, a set of DNA barcode A solutions were introduced to the tissue surface using an array of microchannels for *in situ* ligation of distinct spatial barcode Ai (i = 1-50) to the adapters. Then, a second set of barcodes Bj (j = 1-50) were introduced over the tissue surface in microchannels perpendicularly to those in the first flow barcoding step. They were subsequently ligated at the intersections, resulting in a 2D mosaic of tissue pixels, each of which contains a distinct combination of barcodes Ai and Bj (i = 1-50, j = 1-50). During each flow or afterward, the tissue slides were imaged under an optical microscope such that spatially barcoded accessible chromatin can be correlated with the tissue morphology. After forming a spatially barcoded tissue mosaic (n = 2500), reverse crosslinking was performed to release barcoded DNA fragments, which were amplified by PCR for sequencing library preparation. To evaluate the performance of *in situ* transposition and ligation, the 4′, 6-diamidino-2-phenylindole (DAPI) stained adherent NIH 3T3 cells were fixed by formaldehyde on a glass slide. The cells were then transposed by Tn5 transposase followed by ligation of a dummy barcode A labeled with fluorescein isothiocyanate (FITC) to evaluate the chemistry with fluorescence microscopy. The resulting images revealed a strong overlap between nucleus (blue) and FITC signal (green), indicating the successful insertion of adaptors into accessible chromatin loci with ligated barcode A in nuclei only (Fig. 1b).

**Fig. 1.**
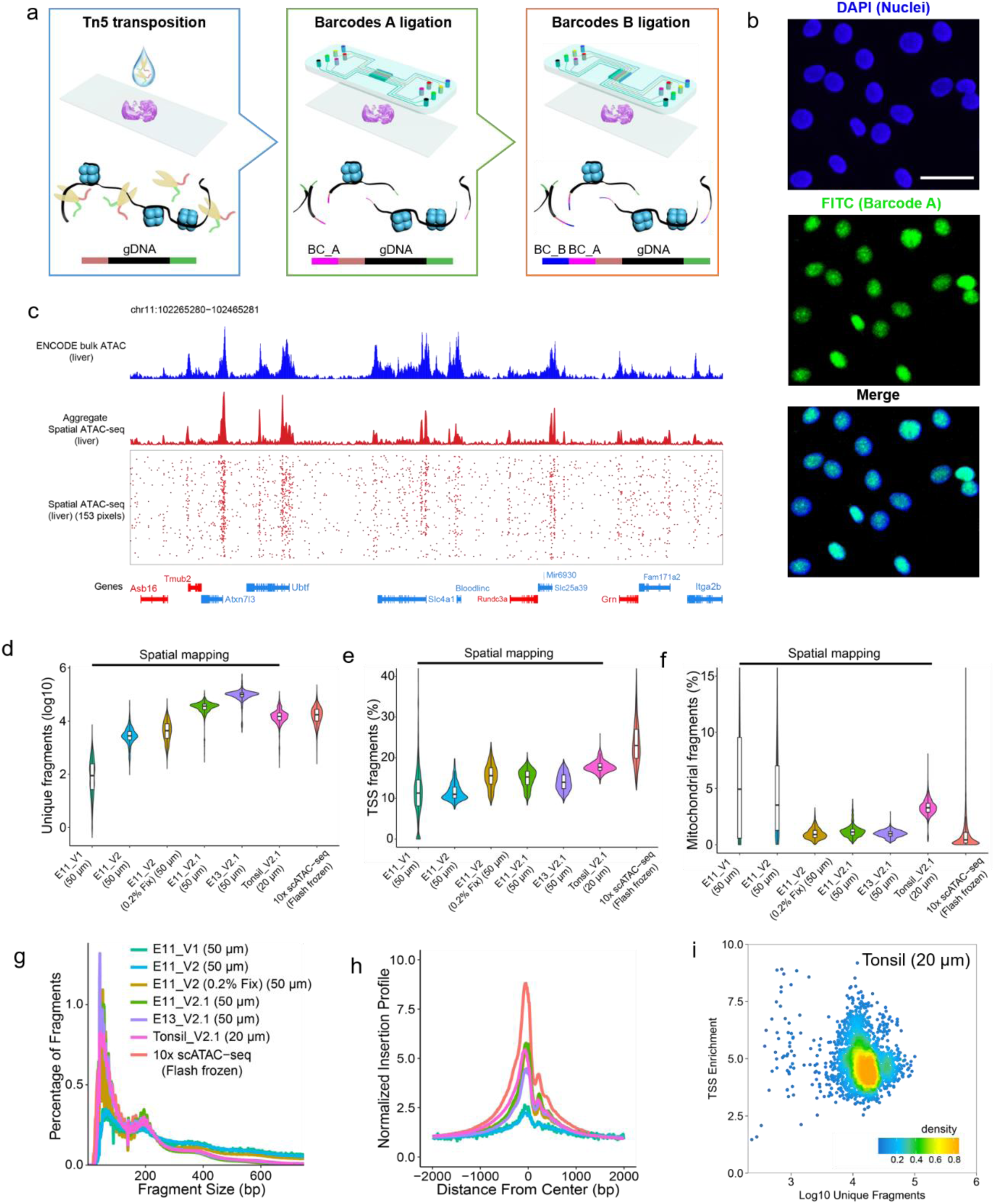
spatial-ATAC-seq: design, workflow, and data quality. **a.** Schematic workflow. Tn5 transposition was performed in tissue sections, followed by in-situ ligation of two sets of DNA barcodes (A1-A50, B1-B50). **b,** Validation of in-situ transposition and ligation using fluorescent DNA probes. Tn5 transposition was performed in 3T3 cells on a glass slide stained by DAPI (blue). Afterwards, FITC-labeled barcode A is ligated to the adapters on the transposase accessible genomic DNA. Scale bar, 50 µm. **c,** Aggregate spatial chromatin accessibility profiles recapitulated published profiles of ATAC-seq in the liver of E13 mouse embryo. **d**, Comparison of number of unique fragments for different protocols and microfluidic channel width between our spatial method in this work and 10x scATAC-seq. **e**, Comparison of fraction of TSS fragments for different protocols and microfluidic channel width between our spatial method in this work and 10x scATAC-seq. **f**, Comparison of fraction of mitochondrial fragments for different protocols and microfluidic channel width between our spatial method in this work and 10x scATAC-seq. **g,** Comparison of insert size distribution of ATAC-seq fragments for different protocols and microfluidic channel width between our spatial method in this work and 10x scATAC-seq. **h,** Comparison of enrichment of ATAC-seq reads around TSSs for different protocols and microfluidic channel width between our spatial method in this work and 10x scATAC-seq. Coloring is consistent with (**g**) **i,** Scatterplot showing the TSS enrichment score vs unique nuclear fragments per cell for human tonsil.

As we proceeded to develop spatial-ATAC-seq, we went through several versions of chemistry to optimize the protocol in order to achieve high yield and high signal-to-noise ratio for the mapping of tissue sections (Fig. 1d-i and Fig. S2a). In chemistry V1, a set of 50 DNA oligomers containing both barcode A and adapter were introduced in microchannels to a tissue section for *in situ* transposition but the efficiency was low due in part to limited amounts of Tn5-DNA in microchannels. In chemistry V2, we conducted bulk transposition followed by two ligation steps to introduce spatial barcodes A-B. We also optimized the fixation condition by reducing formaldehyde concentration from 4% in chemistry V1 to 0.2% in chemistry V2. We tested the sensitivity of different Tn5 transposase enzymes (Diagenode (C01070010) in chemistry V2.1 vs Lucigen (TNP92110) in chemistry V2). The performance measured by the unique fragments detected and the transcription start site (TSS) enrichment score from V1, V2, to V2.1 was summarized in Fig. S2a. We then applied the optimized spatial-ATAC-seq protocol V2.1 to mouse embryos (E11 and E13) and human tonsil, and assessed the data quality by comparison to non-spatial scATAC-seq data from the commercialized platform (10x Genomics). In 50µm spatial ATAC-seq experiments, we obtained a median of 36,303 (E11) and 100,786 (E13) unique fragments per pixel of which 15% (E11) and 14% (E13) of fragments overlap with TSS regions. In addition, proportion of mitochondrial fragments is low for both E11 and E13 (1%). As for the 20µm spatial-ATAC-seq experiment with human tonsil, we obtained a median of 14,939 unique fragments per pixel of which 18% of fragments fell within TSS regions. The fraction of read-pairs mapping to mitochondria is 3%. Overall, the data quality of spatial-ATAC-seq from the tissue section is equivalent to non-spatial scATAC-seq (17,321 unique fragments per cell, 23% TSS fragments, and 0.4% mitochondrial reads). Moreover, the insert size distribution of spatial-ATAC-seq fragments was consistent with the capture of nucleosomal and subnucleosomal fragments for all tissue types (Fig. 1g). We also performed correlation analysis between biological replicates of serial tissue sections for spatial-ATAC-seq, which showed high reproducibility with the Pearson correlation coefficient of 0.95 (Fig. S2b). Using spatial-ATAC-seq, we generated DNA accessibility profiles of individual tissue pixels in the fetal liver of an E13 mouse embryo. Aggregate profiles of spatial ATAC-seq data accurately reproduced the bulk measurement of accessibility obtained from the ENCODE reference database (Fig. 1c).

### Spatial chromatin accessibility mapping of E13 mouse embryo

We next sought to identify cell types *de novo* by chromatin accessibility from the E13 mouse embryo. A pixel by tile matrix was generated by aggregating reads in 5 kilobase bins across the mouse genome. Latent semantic indexing (LSI) and uniform manifold approximation and projection (UMAP) were then applied for dimensionality reduction and embedding, followed by Louvain clustering using the ArchR package^23^. Unsupervised clustering identified 8 main clusters and the spatial map of these clusters revealed distinct patterns that agreed with the tissue histology shown in an adjacent H&E stained tissue section (Fig. 2a to c, Fig. S3). For example, cluster 1 represents the fetal liver in the mouse embryo, and cluster 2 is specific to the spine region, including the dorsal root ganglia (Fig. S4a, b, i, j). Cluster 3 to cluster 5 are associated with the peripheral and central nervous system (PNS and CNS). Cluster 6 includes several cell types present in the developing limbs, and cluster 8 encompasses several developing internal organs. To benchmark spatial-ATAC-seq data, we projected the ENCODE organ-specific ATAC-seq data onto our UMAP embedding using the UMAP transform function^24^. In general, the cluster identification matched well with the bulk ATAC-seq projection (Fig. S3b-d) and distinguished all major developing tissues and organs in a E13 mouse embryo. We further examined cell type-specific marker genes and estimated the expression of these genes from our chromatin accessibility data based on the overall signal at a given locus^23^ (Fig. 2c, Fig. S3e, f). *Sptb*, which plays a role in stability of erythrocyte membranes, was activated extensively in the liver. *Syt8*, which is important in neurotransmission, had a high level of gene activity in the spine. *Ascl1* showed strong enrichment in the mouse brain, which is known to be involved in the commitment and differentiation of neuron and oligodendrocyte (Fig. 2c, Fig. S4e, f). *Sox10* marks oligodendrocyte progenitor cells (OPCs). It was expressed at a high level in the dorsal root ganglia (DRGs), which are adjacent to the spinal cord (Fig. S4a, b). *Olig2* is a marker of neural progenitors, pre-OPCs and OPCs. *Olig2* is expressed in a small domain of the spinal cord, in the ventral domains of the forebrain, and in some posterior regions (brain stem, midbrain and hindbrain), which is consistent with the high gene score in the spatial ATAC-seq data (Fig. S4c, d). However, its expression in forebrain is confined at the dorsal side at this developmental stage as detected by in situ hybridization (Fig. S4c), but the chromatin accessibility is open in both dorsal and ventral side, suggesting the possibility of epigenetic priming. *Ror2* correlates with the early formation of the chondrocytes and cartilage, and it was highly expressed in the limb^25^. Pathway analysis of marker genes revealed that cluster 1 was associated with in erythrocyte differentiation, cluster 5 corresponded to forebrain development, and cluster 6 was involved in limb development, all in agreement with anatomical annotations (Fig. S5). Interestingly, we found that the clusters that appeared to be homogenous could be further deconvoluted into sub-populations with distinct spatial distributions (Fig. S3g). For example, the fetal liver could be further subset to two clusters, and we found that some genes related to hematopoiesis (e.g. *Hbb-y*, *Slc4a1*, *Sptb*) had higher expression in subcluster 1 (Fig. S3g). Moreover, we further investigated the expression patterns in the spine of the E13 mouse embryo and the select genes showing epigenetic gradients along the anterior-posterior axis (Fig. S6).

**Fig. 2.**
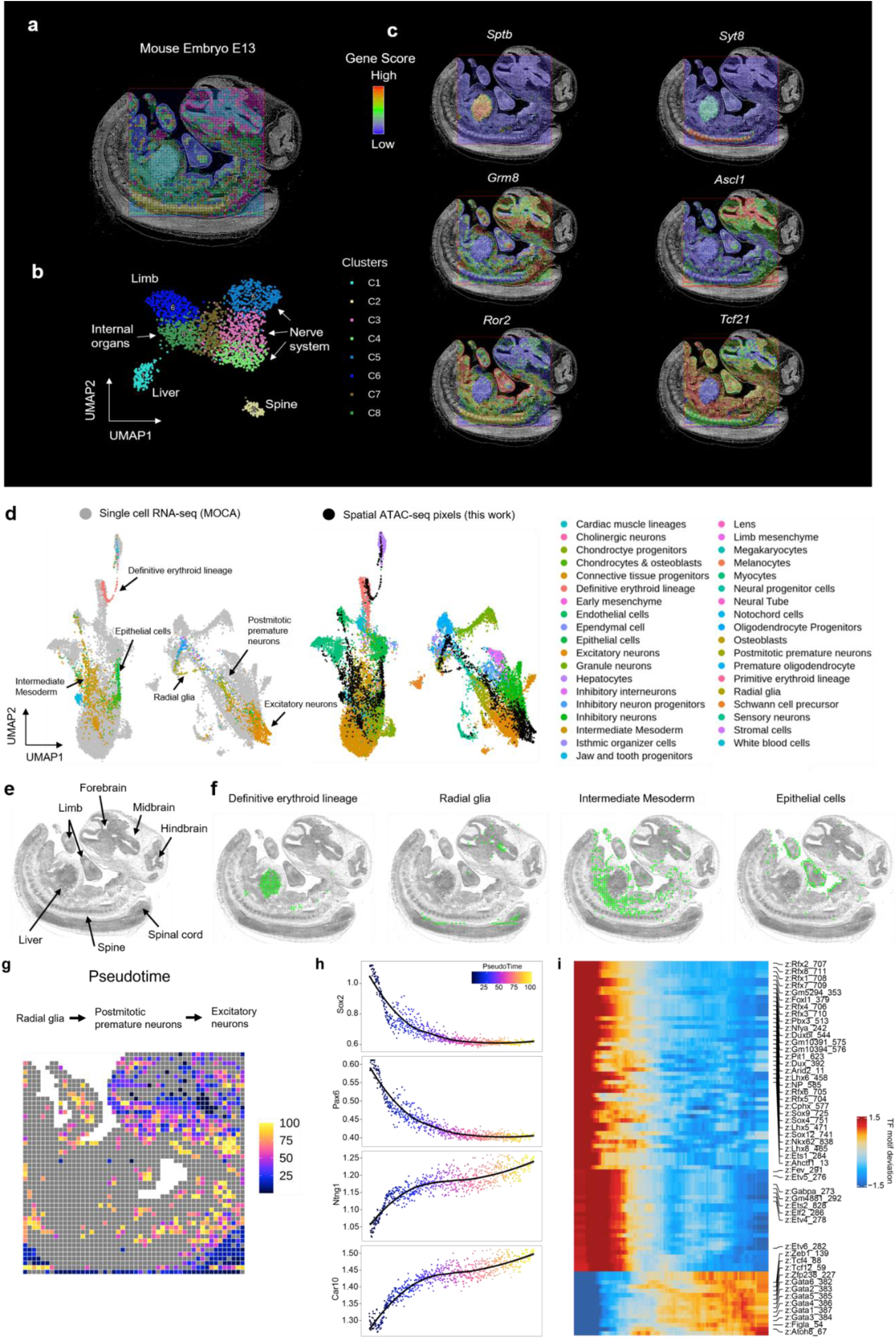
Spatial chromatin accessibility mapping of E13 mouse embryo. **a,** Unbiased clustering analysis was performed based on chromatin accessibility of all tissue pixels (50μm pixel size). Overlay of clusters with the tissue image reveals that the spatial chromatin accessibility clusters precisely match the anatomic regions. **b,** UMAP embedding of unsupervised clustering analysis for chromatin accessibility. Cluster identities and coloring of clusters are consistent with (**a**). **c,** Spatial mapping of gene scores for selected marker genes in different clusters and the chromatin accessibility at select genes are highly tissue specific. **d,** Integration of scRNA-seq from E13.5 mouse embryos^26^ and spatial ATAC-seq data. Unsupervised clustering of the combined data was colored by different cell types. **e,** Anatomic annotation of major tissue regions based on the H&E image. **f,** Spatial mapping of selected cell types identified by label transferring from scRNA-seq to spatial ATAC-seq data. **g,** Pseudotemporal reconstruction from the developmental process from radial glia, postmitotic premature neurons, to excitatory neurons plotted in space. **h,** Dynamics for selected gene score along the pseudo-time shown in (**g**). **i,** Pseudo-time heatmap of TF motifs changes from radial glia, postmitotic premature neurons, to excitatory neurons.

In addition to the inference of cell type-specific marker genes, our approach also enabled the unbiased identification of cell type-specific chromatin regulatory elements (Fig. S7), which provides a resource for defining regulatory elements as cell type-specific reporters. To further utilize the underlying chromatin accessibility data, we sought to examine cell type-specific transcription factor (TF) regulators within each cluster using deviations of TF motifs. We found that the most enriched motifs in the peaks that are more accessible in fetal liver correspond to Gata transcription factors, consistent with their well-studied role in erythroid differentiation (Fig. S7b, c). Cluster 5 enriched for *Sox6* motif that supports its role for the CNS development. *Hoxd11*, which marks the posterior patterning and plays a role in limb morphogenesis, was enriched in the limb (Fig. S7c).

We then integrated the spatial ATAC-seq data with the scRNA-seq data to assign cell types to each cluster^26^ (Fig. 2d-f, Fig. S8a). For example, the definitive erythroid cells were exclusively enriched in the liver. Additionally, we found few hepatocytes and white blood cells in this region, which could not be identified in the E11 data, suggesting that these cell types emerged at the later developmental time points. Intermediate mesoderm was identified in the internal organ region, and radial glia was mainly distributed in the CNS. A refined clustering process also enabled identification of sub-populations in excitatory neurons with distinct spatial distributions, marker genes and chromatin regulatory elements (Fig. S8b-d). During embryonic development, dynamic changes in chromatin accessibility across time and space help regulate the formation of complex tissue architectures and terminally differentiated cell types^27^. In the embryonic CNS, radial glia function as primary progenitors or neural stem cells (NSCs), which give rise to various types of neurons^28^. Therefore, we sought to exploit our spatial ATAC-seq data to recover the spatially organized developmental trajectory and examine how developmental processes proceed across the tissue space. Here, we studied the course of a developmental process from radial glia to excitatory neurons with postmitotic premature neurons as the immediate state after the radial glial differentiation, and ordered these cells in pseudo-time using ArchR. Spatial projection of each pixel’s pseudo-time value revealed the spatially organized developmental trajectory in neurons (Fig. 2g). We then identified changes in gene expression and TF deviations across this developmental process, and many genes recovered are important in neuron development, including *Sox2*, which is required for stem cell maintenance in the central nervous system, and *Ntng1*, which is involved in controlling patterning and neuronal circuit formation (Fig. 2h, i).

### Spatial chromatin accessibility mapping of E11 mouse embryo and comparison with E13 to investigate the spatiotemporal relationship

To map chromatin accessibility during mouse fetal development, we also profiled mouse embryo at E11. Unsupervised clustering identified 4 main clusters with distinct spatial patterns, which showed good agreement with the anatomy in an adjacent H&E stained tissue section (Fig. 3a-c, Fig. S9a-c). Cluster 1 is located in the fetal liver and aorta-gonad-mesonephros (AGM), which are related to embryonic hematopoiesis. It should be noted that spatial ATAC-seq can resolve the fine structure in mouse embryo such as AGM, showing its capability to profile chromatin accessibility in a high spatial resolution manner. Cluster 2 and cluster 3 consist of tissues associated with neuronal development such as mouse brain and neural tube. Cluster 4 includes the embryonic facial prominence, internal organs and limb. In addition, cluster identification matched the ENCODE organ-specific bulk ATAC-seq projection onto the UMAP embedding (Fig. S9d).

**Fig. 3.**
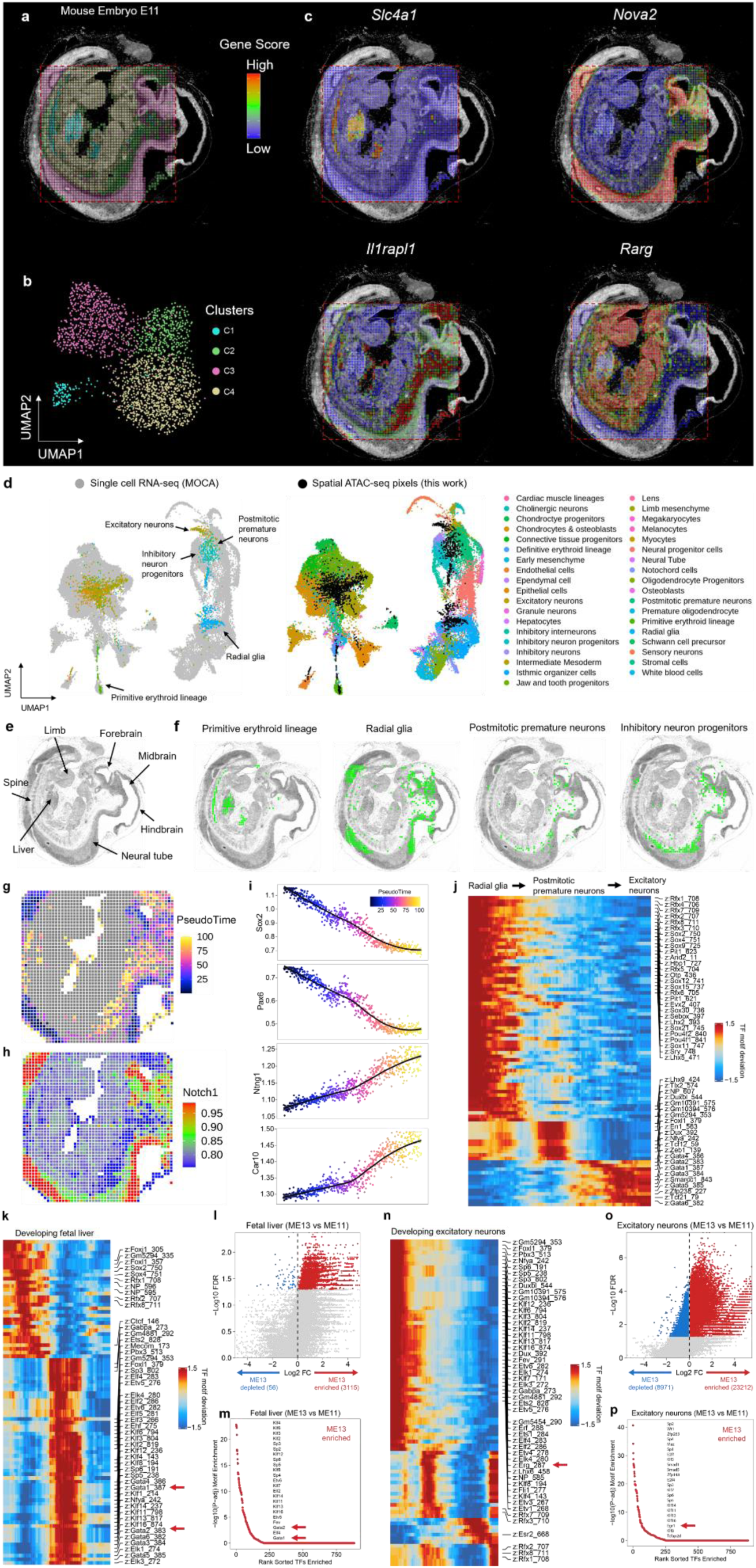
Spatial chromatin accessibility mapping of E11 mouse embryo and spatiotemporal analysis. **a,** Unsupervised clustering analysis and spatial distribution of each cluster. Overlay with the tissue image reveals that the spatial chromatin accessibility clusters precisely match the anatomic regions. **b,** UMAP embedding of unsupervised clustering analysis for chromatin accessibility. Cluster identities and coloring of clusters are consistent with (**a**). **c,** Spatial mapping of gene scores for selected marker genes in different clusters and the chromatin accessibility at select genes are highly tissue specific. **d,** Integration of scRNA-seq from E11.5 mouse embryos^26^ and spatial ATAC-seq data. Unsupervised clustering of the combined data was colored by different cell types. **e,** Anatomic annotation of major tissue regions based on the H&E image. **f,** Spatial mapping of selected cell types identified by label transferring from scRNA-seq to spatial ATAC-seq data. **g,** Pseudotemporal reconstruction from the developmental process from radial glia to excitatory neurons plotted in space. **h,** Spatial mapping of gene scores for *Notch1*. **i,** dynamics for selected gene score along the pseudo-time shown in (**g**). **j,** Pseudo-time heatmap of TF motifs changes from radial glia to excitatory neurons. **k,** Pseudo-time heatmap of TF motifs changes in the fetal liver from E11 to E13 mouse embryo. **l,** Differential peak analysis of fetal liver in E13 mouse embryo compared to E11 mouse embryo. **m,** Ranking of enriched motifs in the peaks that are more accessible in the fetal liver of E13 mouse embryo compared to E11 mouse embryo. **n,** Pseudo-time heatmap of TF motifs changes in the excitatory neurons from E11 to E13 mouse embryo. **o,** Differential peak analysis of excitatory neurons in E13 mouse embryo compared to E11 mouse embryo. **p,** Ranking of enriched motifs in the peaks that are more accessible in the excitatory neurons of E13 mouse embryo compared to E11 mouse embryo.

We further surveyed the chromatin accessibility patterns that distinguished each cluster (Fig. 3c, Fig. S9e, f). For example, *Slc4a1*, which are required for normal flexibility and stability of the erythrocyte membrane and for normal erythrocyte shape, were highly active in liver and AGM. *Nova2*, which is involved in RNA splicing or metabolism regulation in a specific subset of developing neurons, was highly enriched in the brain and neural tube. *Rarg*, which plays an essential role in limb bud development, skeletal growth, and matrix homeostasis, was activated extensively in the embryonic facial prominence and limb^25^. Moreover, we conducted Gene Ontology (GO) enrichment analysis for each cluster, and the GO pathways identified the development processes consistent with the anatomical annotation (Fig. S10). To gain deeper insights into the regulatory factors in each tissue, we clustered chromatin regulatory elements and examined enrichment for TF binding motifs, and expression patterns of those motifs (Fig. S11). We observed strong enrichment of the motifs for *Gata2* (Fig. S10b) and *Ascl2* (Fig. S11c) in the clusters associated with embryonic hematopoiesis and neuronal development, respectively. These master regulators further validated the unique identity of each cluster.

To assign cell types to each cluster, we integrated the spatial ATAC-seq data with the scRNA-seq atlas of the mouse embryos^26^, and several organ-specific cell types were identified (Fig. 3d-f, Fig. S12). The primitive erythroid cells, crucial for early embryonic erythroid development, were strongly enriched in the liver and AGM in agreement with the anatomical annotation. Radial glia, postmitotic premature neurons, and inhibitory neuron progenitors were found in the brain and neural tube. Compared to E13, higher proportion of radial glia were identified in E11 mouse embryo, suggesting their transient nature during CNS development^29^. We observed abundant chondrocytes & osteoblasts in the embryonic facial prominence, and the limb mesenchyme was highly enriched in the limb region. We also reconstructed the spatially organized neuronal development trajectory from radial glia to excitatory neurons in E11 mouse embryo (Fig. 3g-j) and identified the changes in neuron development-related genes and TF deviations across this developmental process, including *Notch1* that is highly expressed in the radial glia and regulates neural stem cell number and function during development^29,30^ (Fig. 3h).

To assess the temporal dynamics of chromatin accessibility more directly during development, we identified dynamic peaks that exhibit a significant change in accessibility from E11 to E13 mouse embryo within fetal liver and excitatory neurons. We observed significant differences in the chromatin accessibility of fetal liver and excitatory neurons between different developmental stages (Fig. 3k-p). In particular, chromatin accessibility profiles of fetal liver at E13 were enriched with Gata motif sequences (Fig. 3k, m), the TFs known to be important in the erythroid differentiation^23^. In addition, *Egr1* motif was enriched in the excitatory neurons at E13, which has the functional implication during brain development, particularly for the specification of excitatory neurons^31^.

### Spatial chromatin accessibility mapping of human tonsil and immune cell states

To demonstrate the ability to profile spatial chromatin accessibility in different tissue types and species, we then applied spatial-ATAC-seq with 20 µm pixel size to the human tonsil tissue. Unsupervised clustering revealed distinct spatial features with the germinal centers (GC) identified mainly in cluster 1 (Fig. 4a-c). We set out to explore the spatial patterns of specific marker genes to distinguish cell types (Fig. 4d, Fig. S14) and compared to the distribution of protein expression in tonsil (Fig. S15). For B cell-related genes, the accessibility of *CD10*, a marker for mature GC B cells, was enriched in the GC regions. *CD27*, a marker for memory B cells, was active in GC and the extrafollicular regions. *CD38*, which marks activated B cells, was found to be enriched in GC. *CXCR4*, which is expressed in the centroblasts in the GC dark zone, unexpectedly showed high accessibility only in non-GC cells. This discordance between epigenetic state and protein expression may suggest epigenetic priming of pre-GC B cells prior to entering GC. It could also be due to the presence of CXCR4+ T cells supporting extra-follicular B-cell responses in the setting of inflammation^32^. *PAX5*, a transcription factor for follicular and memory B cells, was enriched in GC but also observed in the extrafollicular zones where the memory B cells migrated to. *BHLHE40*, a poorly understood transcription factor that can bind to the major regulatory regions of the IgH locus, was found to be enriched in the extrafollicular region but completely depleted in GC, suggesting the potential role in the regulation of class switch recombination in the pre-GC state. This supports a model of epigenetic control for class switch recombination that occurs before formation of the GC response. For T cell related genes, *CD3* corresponded to T cell zones and also found active in GC. It is known that follicular helper T cells (T_FH_) trafficking into GC requires downregulation of CCR7 and upregulation of CXCR5. We observed significantly reduced *CCR7* accessibility in GC while strong enrichment outside GC, indicating this T_FH_ function is indeed epigenetically regulated. *CXCR5* accessibility was extensively detected in GC but also observed outside GC, indicating a possible early epigenetic priming of T_FH_ cells prior to GC entry for B cell help. The locus accessibility of *BCL6*, a T_FH_ master transcription factor, was strongly enriched in GC as expected. *FOXP3*, a master transcription factor for follicular regulatory T cells (T_FR_), is mainly in the extrafollicular zone but at low frequency according to human protein atlas data (Fig. S15). Interestingly, it showed extensive open locus accessibility, suggesting extensive epigenetic priming of pre-GC T cells to potentially develop T_FR_ function as needed to balance GC activity. *CD25*, a surface marker for regulatory T cells, was active in both GC and the extrafollicular zone. For non-lymphoid cells, *CD11B*, a macrophage marker, was inactive in GC, on contrast to *CD11A,* which was more active in GC lymphocytes. *CD103* was enriched in GC follicular dendritic cells. *CD144*, which encodes vascular endothelial cadherin (VE-cadherin), corresponded to endothelial microvasculature near the crypt or between follicles. *CD32*, a surface receptor involved in phagocytosis and clearing of immune complexes, and *CD55*, a complement decay-accelerating factor, were both active in the same region such that the cells not supposed to be cleared can be protected against phagocytosis by blocking the formation of the membrane attack complex. We also examined cell type-specific TF regulators within each cluster and our data revealed that KLF family transcription factors were highly enriched in non-germinal center cells, consistent with previous study^33^ (Fig. S16).

**Fig. 4.**
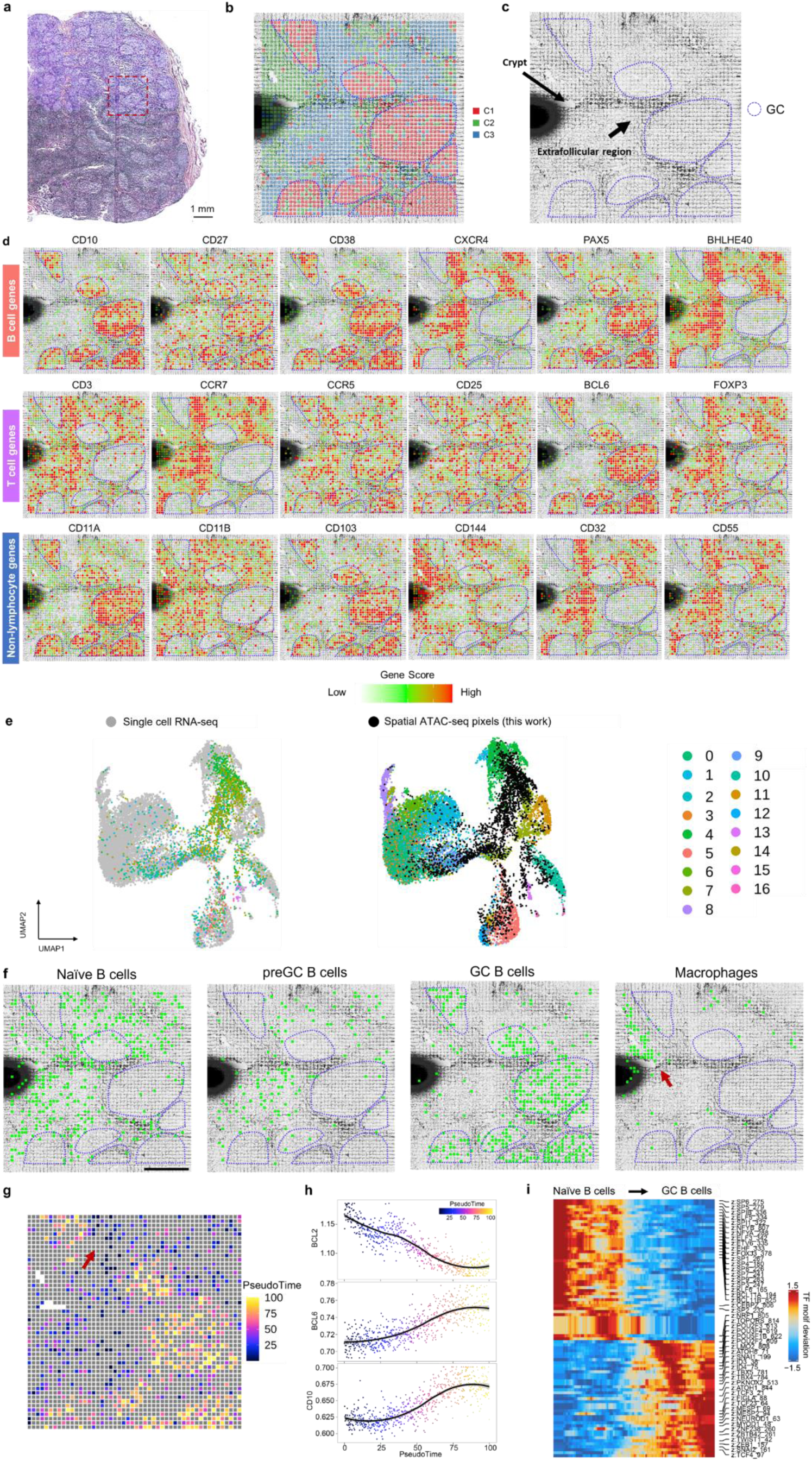
Spatial chromatin accessibility mapping of human tonsil with 20 µm pixel size. **a,** H&E image of a human tonsil from an adjacent tissue section and a region of interest for spatial chromatin accessibility mapping. **b,** Unsupervised clustering analysis and spatial distribution of each cluster. **c,** Anatomic annotation of major tonsillar regions. **d,** Spatial mapping of gene scores for selected genes. **e,** Integration of scRNA-seq data^33^ and spatial ATAC-seq data. Unsupervised clustering of the combined data was colored by different cell types. **f,** Spatial mapping of selected cell types identified by label transferring from scRNA-seq to spatial ATAC-seq data. Scale bar, 500 µm. **g,** Pseudotemporal reconstruction from the developmental process from Naïve B cells to GC B cells plotted in space. **h,** dynamics for selected gene score along the pseudo-time shown in (**g**). **i,** Pseudo-time heatmap of TF motifs changes from Naïve B cells to GC B cells.

To map cell types onto each cluster, we integrated our spatial-ATAC-seq data with the publicly available tonsillar scRNA-seq datasets^33^. After unsupervised clustering for scRNA-seq data and label transfer to the spatial-ATAC-seq data, we found that cells from cluster 0 were widely distributed in the non-GC region, while cells from cluster 4 were enriched in GC (Fig. 4e, f, Fig. S13a). We also identified a small region with cells enriched from cluster 13 (Fig. 4f, Fig. S13a). To define the cell identities for scRNA-seq clusters, we examined the marker genes for each cluster and found that cluster 0 comprised of Naïve B cells, cluster 4 corresponded to GC B cells, and cluster 13 were macrophages (Fig. S13b), in agreement with the tissue histology (Fig. 4f).

Lymphocyte activation, maturation, and differentiation are regulated by the gene networks under the control of transcription factors^33^. To understand the dynamic regulation process, we implemented a pseudotemporal reconstruction of B cell activation to the GC reaction (Fig. 4g-i). Meanwhile, the projection of each pixel’s pseudo-time value onto spatial coordinates revealed spatially distinct regions in this dynamic process. Interestingly, we found that the enriched macrophage population was co-localized with inactivated B cell, consistent with the fact that B cells are activated through acquiring antigen from the antigen presenting macrophages before GC entry or formation^34^ (Fig. 4g). In addition, pseudotemporal ordering of B cell activation revealed dynamic expression and chromatin activity before commitment to the GC state (Fig. 4h, i), including an early activity of BCL2 and reduced accessibility within GC B cells as compared to naïve populations, suggesting that this anti-apoptotic molecule may be actively repressed to ensure that GC B cells are eliminated by apoptosis if they are not selected and rescued by survival signals. In contrast, *LMO2* exhibited increased accessibility at the target sites within GC B cells, which agreed with the previous finding that *LMO2* is specifically upregulated in the GC^35^.

## Discussion

We developed spatial-ATAC-seq for spatially resolved unbiased and genome-wide profiling of chromatin accessibility in intact tissue sections with the pixel size (20μm) at cellular level. The data quality was excellent with ∼15,000 unique fragments detected per 20µm pixel and up to ∼100,000 unique fragments per 50µm pixel. It was applied to mouse embryos (E11 and E13) to delineate the epigenetic landscape of organogenesis, identified all major tissue types with distinct chromatin accessibility state, and revealed the spatiotemporal changes in development. It was also applied to mapping the epigenetic state of different immune cells in human tonsil and revealed the dynamics of B cell activation to GC reaction. The limitations or the areas for further development include the following. First, seamless integration with high-resolution tissue images, i.e., multicolor immunofluorescence image, to identify the cells in each pixel. We observed that a significant number of pixels (20μm) contained single nuclei and the extraction of sequencing reads from these pixels can give rise to spatially-defined single-cell ATAC-seq data. Second, integration with other spatial omics measurements such as transcriptome and proteins, to provide a comprehensive picture of cell types and cell states within the spatial context of tissue. We may simply combine reagents for DBiT-seq^14^ and spatial-ATAC-seq in the same microfluidic barcoding step to achieve spatial multi-omics profiling, which should work in theory but does require further optimization for tissue fixation and reaction conditions to make these assays compatible. Third, it is yet to be further extended to human disease tissues to realize the full potential of spatial-ATAC-seq in clinical research. We anticipate that spatial-ATAC-seq will add a new dimension to spatial biology, which may transform multiple biomedical research fields including developmental biology, neuroscience, immunology, oncology, and clinical pathology, thus empowering scientific discovery and translational medicine in human health and disease.

## METHODS

### Fabrication and assembly of microfluidic device

The molds for microfluidic devices were fabricated in the cleanroom with standard photo lithography. We followed the manufacturer’s guidelines to spin coat SU-8 negative photoresist (SU-2010, SU-2025, Microchem) on a silicon wafer (C04004, WaferPro). The feature heights of 50-µm-wide and 20-µm-wide microfluidic channel device were about 50 µm and 23 µm, respectively. During UV light exposure, chrome photomasks (Front Range Photomasks) were used. Soft lithography was used for polydimethylsiloxane (PDMS) microfluidic devices fabrication. We mixed base and curing agent at a 10:1 ratio and added it over the SU-8 masters. The PDMS was cured (65 °C, 2 h) after degassing in vacuum (30 min). After solidification, PDMS slab was cut out. The outlet and inlet holes were punched for further use.

### Preparation of tissue slides

Mouse C57 Embryo Sagittal Frozen Sections (MF-104-11-C57) and Human Tonsil Frozen Sections (HF-707) were purchased from Zyagen (San Diego, CA). Tissues were snapped frozen in OCT (optimal cutting temperature) compounds, sectioned (thickness of 7-10 µm) and put at the center of poly-L-lysine covered glass slides (63478-AS, Electron Microscopy Sciences).

### H&E staining

The frozen slide was warmed at room temperature for 10 min and fixed with 1mL 4% formaldehyde (10 min). After being washed once with 1X DPBS, the slide was quickly dipped in water and dried with air. Isopropanol (500 μl) was then added to the slide and incubate for 1 minute before being removed. After completely dry in the air, the tissue section was stained with 1 mL hematoxylin (Sigma) for 7 min and cleaned in DI water. The slide was then incubated in 1 mL bluing reagent (0.3% acid alcohol, Sigma) for 2 min and rinsed in DI water. Finally, the tissue slide was stained with 1 mL eosin (Sigma) for 2 min and cleaned in DI water.

### Preparation of transposome

Unloaded Tn5 transposase (C01070010) was purchased from Diagenode, and the transposome was assembled following manufacturer’s guidelines. The oligos used for transposome assembly were as follows:

Tn5MErev:

5′-/5Phos/CTGTCTCTTATACACATCT-3′

Tn5ME-A:

5′-/5Phos/CATCGGCGTACGACTAGATGTGTATAAGAGACAG-3′

Tn5ME-B:

5′-GTCTCGTGGGCTCGGAGATGTGTATAAGAGACAG-3′

### DNA oligos, DNA barcodes sequences, and other key reagents

DNA oligos used for sequencing library construction and PCR were listed in Table S1, DNA barcodes sequences were shown in Table S2, and all other key reagents were given in Table S3.

### Spatial ATAC-seq profiling

The frozen slide was warmed at room temperature for 10 min. The tissue was fixed with formaldehyde (0.2%, 5 min) and quenched with glycine (1.25 M, 5 min) at room temperature. After fixation, the tissue was washed twice with 1 mL 1X DPBS and cleaned in DI water. The tissue section was then permeabilized with 500 µL lysis buffer (10 mM Tris-HCl, pH 7.4; 10 mM NaCl; 3 mM MgCl_2_; 0.01% Tween-20; 0.01% NP-40; 0.001% Digitonin; 1% BSA) for 15 min and was washed by 500 µL wash buffer (10 mM Tris-HCl pH 7.4; 10 mM NaCl; 3 mM MgCl_2_; 1% BSA; 0.1% Tween-20) for 5 min. 100 µL transposition mix (50 µL 2X tagmentation buffer; 33 µL 1X DPBS; 1 µL 10% Tween-20; 1 µL 1% Digitonin; 5 µL transposome; 10 µL Nuclease-free H_2_O) was added followed by incubation at 37 °C for 30 min. After removing transposition mix, 500 µL 40 mM EDTA was added for incubation at room temperature for 5 min to stop transposition. Finally, the EDTA was removed, and the tissue section was washed with 500 µL 1X NEBuffer 3.1 for 5 min.

For barcodes A in situ ligation, the 1st PDMS slab was used to cover the region of interest, the brightfield image was taken with 10X objective (Thermo Fisher EVOS fl microscope) for further alignment. The tissue slide and PDMS device were then clamped with an acrylic clamp. First, DNA barcodes A was annealed with ligation linker 1, 10 μL of each DNA Barcode A (100 μM), 10 μL of ligation linker (100 μM) and 20 μL of 2X annealing buffer (20 mM Tris, pH 7.5-8.0, 100 mM NaCl, 2 mM EDTA) were added together and mixed well. Then, 5 μL ligation reaction solution (50 tubes) was prepared by adding 2 μL of ligation mix (72.4 μL of RNase free water, 27 μL of T4 DNA ligase buffer, 11 μL T4 DNA ligase, 5.4 μL of 5% Triton X-100), 2 μL of 1X NEBuffer 3.1 and 1 μL of each annealed DNA barcode A (A1-A50, 25 μM) and loaded into each of the 50 channels with vacuum. The chip was kept in a wet box for incubation (37 °C, 30 min). After flowing through 1X NEBuffer 3.1 for washing (5 min), the clamp and PDMS were removed. The slide was quickly dipped in water and dried with air.

For barcodes B in situ ligation, the 2nd PDMS slab with channels perpendicular to the 1st PDMS was attached to the dried slide carefully. A brightfield image was taken and the acrylic clamp was used to press the PDMS against the tissue. The annealing of DNA barcodes B with ligation linker 2 were the same with DNA barcodes A and ligation linker 1 annealing. The preparation and addition of ligation reaction solution for DNA barcode B (B1-B50, 25 μM) were also the same with DNA barcode A (A1-A50, 25 μM). The chip was kept in a wet box for incubation (37 °C, 30 min). After flowing through 1X DPBS for washing (5 min), the clamp and PDMS were removed, the tissue section was dipped in water and dried with air. The final brightfield image of the tissue was taken.

For tissue digestion, the interest region of the tissue was covered with a square PDMS well gasket and 100 μL reverse crosslinking solution (50 mM Tris-HCl, pH 8.0; 1 mM EDTA; 1% SDS; 200 mM NaCl; 0.4 mg/mL proteinase K) was loaded into it. The lysis was conducted in a wet box (58 °C, 2 h). The final tissue lysate was collected into a 200 μL PCR tube for incubation with rotation (65 °C, overnight).

For library construction, the lysate was first purified with Zymo DNA Clean & Concentrator-5 and eluted to 20 μL of DNA elution buffer, followed by mixing with the PCR solution (2.5 µL 25 µM new P5 PCR primer; 2.5 µL 25 µM Ad2 primer; 25 µL 2x NEBNext Master Mix). Then, PCR was conducted with following the program: 72 °C for 5 min, 98 °C for 30 s, and then cycled 5 times at 98 °C for 10 s, 63 °C for 10 s, and 72°C for 1 min. To determine additional cycles, 5 µL of the pre-amplified mixture was first mixed with the qPCR solution (0.5 µL 25 µM new P5 PCR primer; 0.5 µL 25 µM Ad2 primer; 0.24 µl 25x SYBR Green; 5 µL 2x NEBNext Master Mix; 3.76 µL nuclease-free H_2_O). Then, qPCR reaction was carried out at the following conditions: 98 °C for 30 s, and then 20 cycles at 98 °C for 10 s, 63 °C for 10 s, and 72°C for 1 min. Finally, the remainder 45 µL of the pre-amplified DNA was amplified by running the required number of additional cycles of PCR (cycles needed to reach 1/3 of saturated signal in qPCR).

To remove PCR primers residues, the final PCR product was purified by 1X Ampure XP beads (45 µL) following the standard protocol and eluted in 20 µL nuclease-free H_2_O. Before sequencing, an Agilent Bioanalyzer High Sensitivity Chip was used to quantify the concentration and size distribution of the library. Next Generation Sequencing (NGS) was performed using the Illumina HiSeq 4000 sequencer (pair-end 150 bp mode with custom read 1 primer).

### Data preprocessing

Two constant linker sequences (linker 1 and linker 2) were used to filter Read 1, and the filtered sequences were transformed to Cell Ranger ATAC format (10x Genomics). The genome sequences were in the new Read 1, barcodes A and barcodes B were included in new Read 2. Resulting fastq files were aligned to the mouse reference (mm10) or human reference (GRCh38), filtered to remove duplicates and counted using Cell Ranger ATAC v1.2. The BED like fragments file were generated for downstream analysis. The fragments file contains fragments information on the genome and tissue location (barcode A x barcode B). A preprocessing pipeline we developed using Snakemake workflow management system is shared at https://github.com/dyxmvp/Spatial_ATAC-seq.

### Data visualization

We first identified pixels on tissue with manual selection from microscope image using Adobe Illustrator (https://github.com/rongfan8/DBiT-seq), and a custom python script was used to generate metadata files that were compatible with Seurat workflow for spatial datasets.

The fragment file was read into ArchR as a tile matrix with the genome binning size of 5kb, and pixels not on tissue were removed based on the metadata file generated from the previous step. Data normalization and dimensionality reduction was conducted using iterative Latent Semantic Indexing (LSI) (iterations = 2, resolution = 0.2, varFeatures = 25000, dimsToUse = 1:30, sampleCells = 10000, n.start = 10), followed by graph clustering and Uniform Manifold Approximation and Projection (UMAP) embeddings (nNeighbors = 30, metric = cosine, minDist = 0.5)^36^.

Gene Score model in ArchR was employed to gene accessibility score. Gene Score Matrix was generated for downstream analysis. The getMarkerFeatures and getMarkers function in ArchR (testMethod = “wilcoxon”, cutOff = “FDR <= 0.05 & Log2FC >= 0.25”) was used to identify the marker regions/genes for each cluster, and gene scores imputation was implemented with addImputeWeights for data visualization. The enrichGO function in the clusterProfiler package was used for GO enrichment analysis (qvalueCutoff = 0.05)^37^. For spatial data visualization, results obtained in ArchR were loaded to Seurat V3.2.3 to map the data back to the tissue section^38,39^.

In order to project bulk ATAC-seq data, we downloaded raw sequence data aligned to mm10 (BAM files) from ENCODE. After counting the reads in 5kb tiled genomes using getCounts function in chromVAR^40^, the projectBulkATAC function in ArchR was used.

Cell type identification and pseudo-scRNA-seq profiles was added through integration with scRNA-seq reference data^26^. FindTransferAnchors function (Seurat V3.2 package) was used to align pixels from spatial ATAC-seq with cells from scRNA-seq by comparing the spatial ATAC-seq gene score matrix with the scRNA-seq gene expression matrix. GeneIntegrationMatrix function in ArchR was used to add cell identities and pseudo-scRNA-seq profiles.

Pseudobulk group coverages based on cluster identities were generated with addGroupCoverages and used for peak calling with macs2 using addReproduciblePeakSet function in ArchR. To compute per-cell motif activity, chromVAR^40^ was run with addDeviationsMatrix using the cisbp motif set after a background peak set was generated using addBgdPeaks. Cell type-specific marker peaks were identified with getMarkerFeatures (bias = c(”TSSEnrichment”, “log10(nFrags)”), testMethod = “wilcoxon”) and getMarkers (cutOff = “FDR <= 0.05 & Log2FC >= 0.1”). Pseudotemporal reconstruction was implemented by addTrajectory function in ArchR.

### Published data for data quality comparison and integrative data analysis

10x scATAC-seq (Flash frozen): Flash frozen cortex, hippocampus, and ventricular zone from embryonic mouse brain (E18). (Single Cell ATAC Dataset by Cell Ranger ATAC 1.2.0)

ENCODE (bulk): Public bulk ATAC-seq datasets were downloaded from ENCODE (E11.5 and E13.5).

Mouse organogenesis cell atlas (MOCA): https://oncoscape.v3.sttrcancer.org/atlas.gs.washington.edu.mouse.rna/downloads

Human tonsillar scRNA-seq: Gene Expression Omnibus under accession GSE165860.

## Reporting summary

Further information on research design is available in the Nature Research Reporting Summary linked to this paper.

## Code availability

Code for sequencing data analysis is available on Github: https://github.com/dyxmvp/Spatial_ATAC-seq.

## Data availability

Raw and processed data reported in this paper are deposited in the Gene Expression Omnibus (GEO) with accession code GSE171943.

## Acknowledgments

We thank the Yale Center for Research Computing for guidance and use of the research computing infrastructure. The molds for microfluidic devices were fabricated at the Yale University School of Engineering and Applied Science (SEAS) Nanofabrication Center. Next-generation sequencing was conducted at Yale Stem Cell Center Genomics Core Facility which was supported by the Connecticut Regenerative Medicine Research Fund and the Li Ka Shing Foundation. Service provided by the Genomics Core of Yale Cooperative Center of Excellence in Hematology (U54DK106857) was used. This research was supported by Packard Fellowship for Science and Engineering (to R.F.), Stand-Up-to-Cancer (SU2C) Convergence 2.0 Award (to R.F.), and Yale Stem Cell Center Chen Innovation Award (to R.F.). It was supported in part by grants from the U.S. National Institutes of Health (NIH) (U54CA209992, R01CA245313, and UG3CA257393, to R.F.). Y.L. was supported by the Society for ImmunoTherapy of Cancer (SITC) Fellowship.

## Contributions

Conceptualization: R.F.; Methodology: Y.D., D.Z., and Y.L.; Experimental Investigation: Y.D. and D.Z.; Data Analysis: Y.D., G.C.-B., and R.F.; Resources: X.Q. and G.S.; M.B., S.M., M.L.X., S.H., and J.E.C. provided valuable advice and input; Original Draft: Y.D., D.Z., and R.F. All authors reviewed, edited, and approved the manuscript.

## Competing interests

R.F. and Y.D. are inventors of a patent application related to this work. R.F. is scientific founder and advisor of IsoPlexis, Singleron Biotechnologies, and AtlasXomics. The interests of R.F. were reviewed and managed by Yale University Provost’s Office in accordance with the University’s conflict of interest policies.

---

**Fig. S1.**
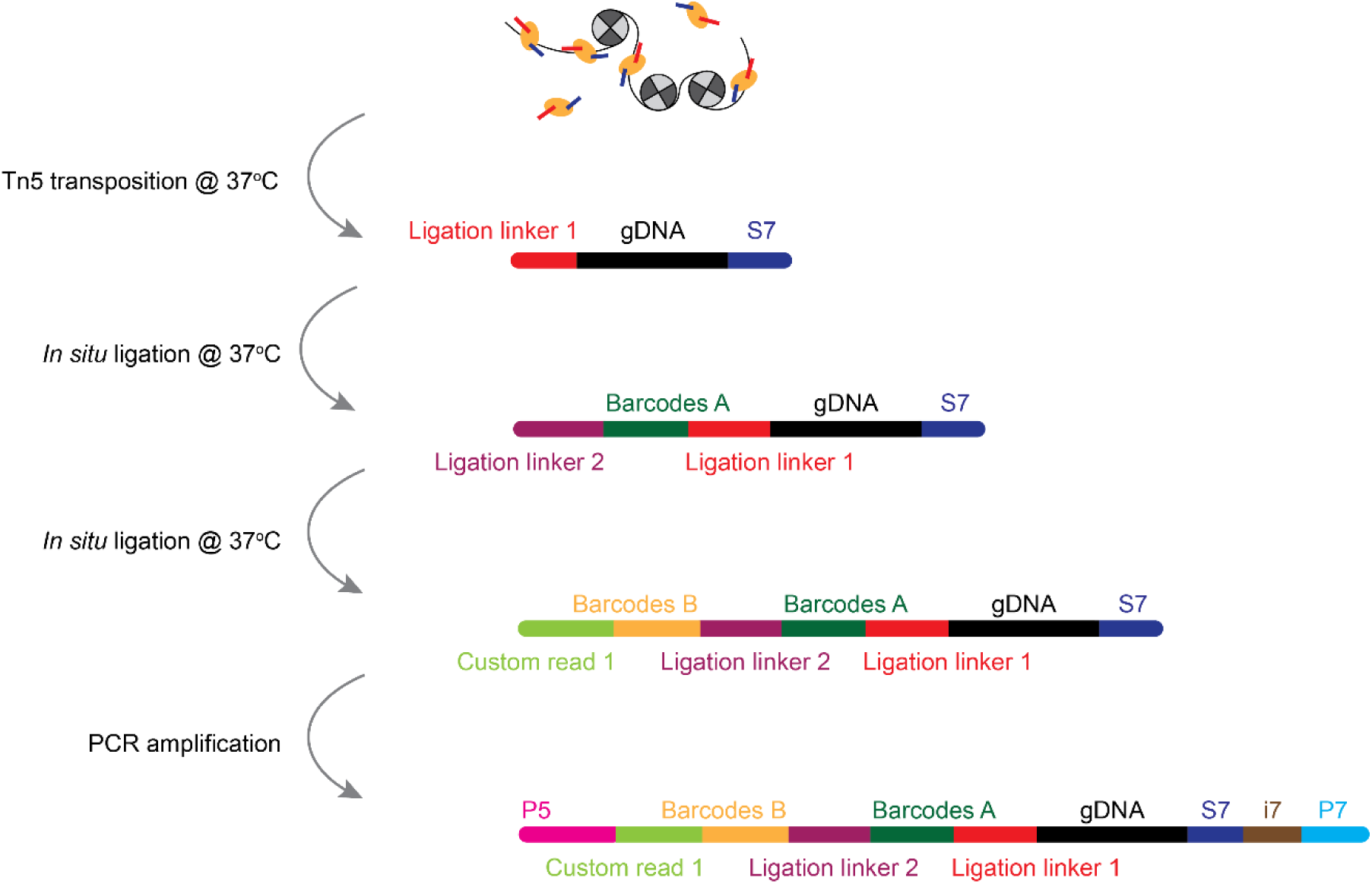
Chemistry workflow of spatial-ATAC-seq. A tissue section on a standard aminated glass slide was lightly fixed with formaldehyde. Then, Tn5 transposition was performed at 37 °C, and the adapters containing ligation linker 1 were inserted to the cleaved genomic DNA at transposase accessible sites. Afterwards, a set of DNA barcode A solutions were introduced by microchannel-guided flow delivery to perform in situ ligation reaction for appending a distinct spatial barcode Ai (i = 1-50) and ligation linker 2. Then, a second set of barcodes Bj (j = 1-50) were introduced using another set of microfluidic channels perpendicularly to those in the first flow barcoding step, which were subsequently ligated at the intersections, resulting in a mosaic of tissue pixels, each containing a distinct combination of barcodes Ai and Bj (i = 1-50, j = 1-50). After DNA fragments were collected by reversing cross-linking, the library construction was completed during PCR amplification.

**Fig. S2.**
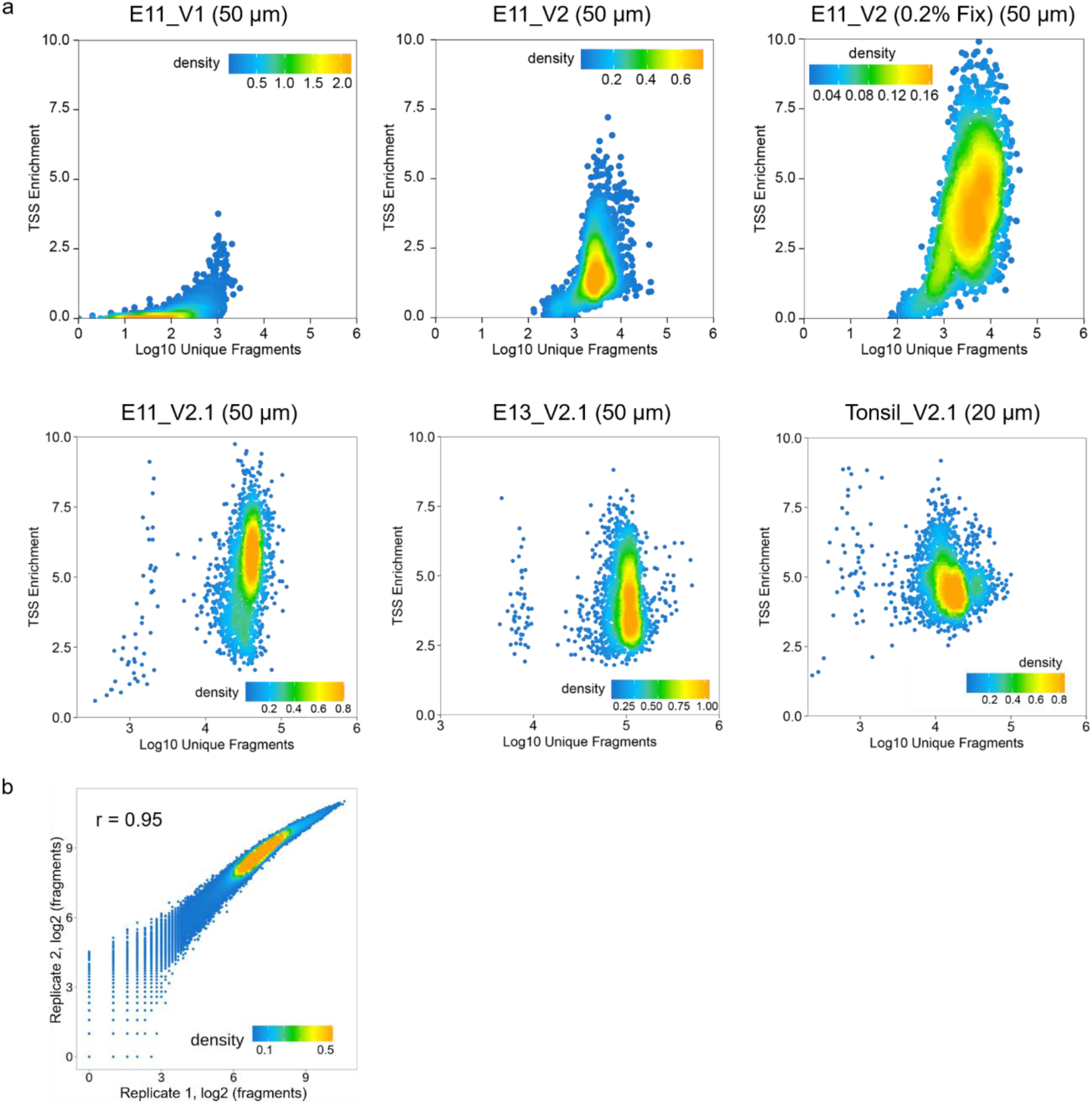
Quality control metrics for spatial ATAC-seq datasets. **a,** Scatterplot showing the TSS enrichment score vs unique nuclear fragments per cell for different protocols and microfluidic channel width. **b,** Reproducibility between biological replicates on E13 mouse embryo. Pearson correlation coefficient r = 0.95.

**Fig. S3.**
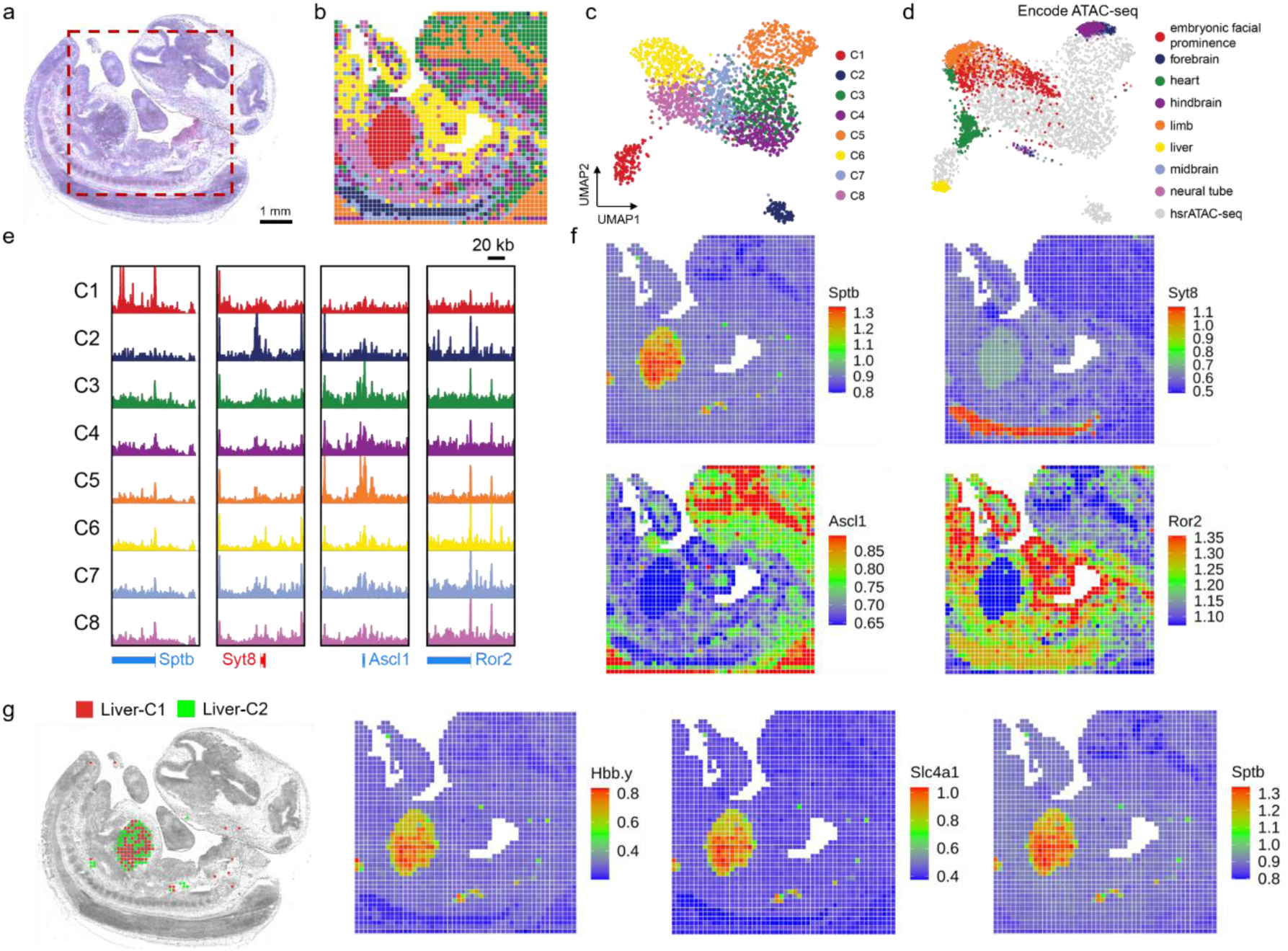
Further analysis of spatial chromatin accessibility mapping of E13 mouse embryo, validation with ENCODE, and sub-clustering in liver. **a,** H&E image from an adjacent tissue section and a region of interest for spatial chromatin accessibility mapping (50 µm pixel size). **b,** Unsupervised clustering analysis and spatial distribution of each cluster. **c,** UMAP embedding of unsupervised clustering analysis for spatial ATAC-seq. Cluster identities and coloring of clusters are consistent with (**b**). **d,** LSI projection of ENCODE bulk ATAC-seq data from diverse cell types of the E13.5 mouse embryo dataset onto the spatial ATAC-seq embedding. **e, f,** Genome browser tracks (**e**) and spatial mapping (**f**) of gene scores for selected marker genes in different clusters. **g,** Refined clustering of fetal liver in E13 mouse embryo enabled identification of sub-populations, and some genes related to hematopoiesis (e.g. *Hbb-y*, *Slc4a1*, *Sptb*) had higher expression lever in the subcluster 1.

**Fig. S4.**
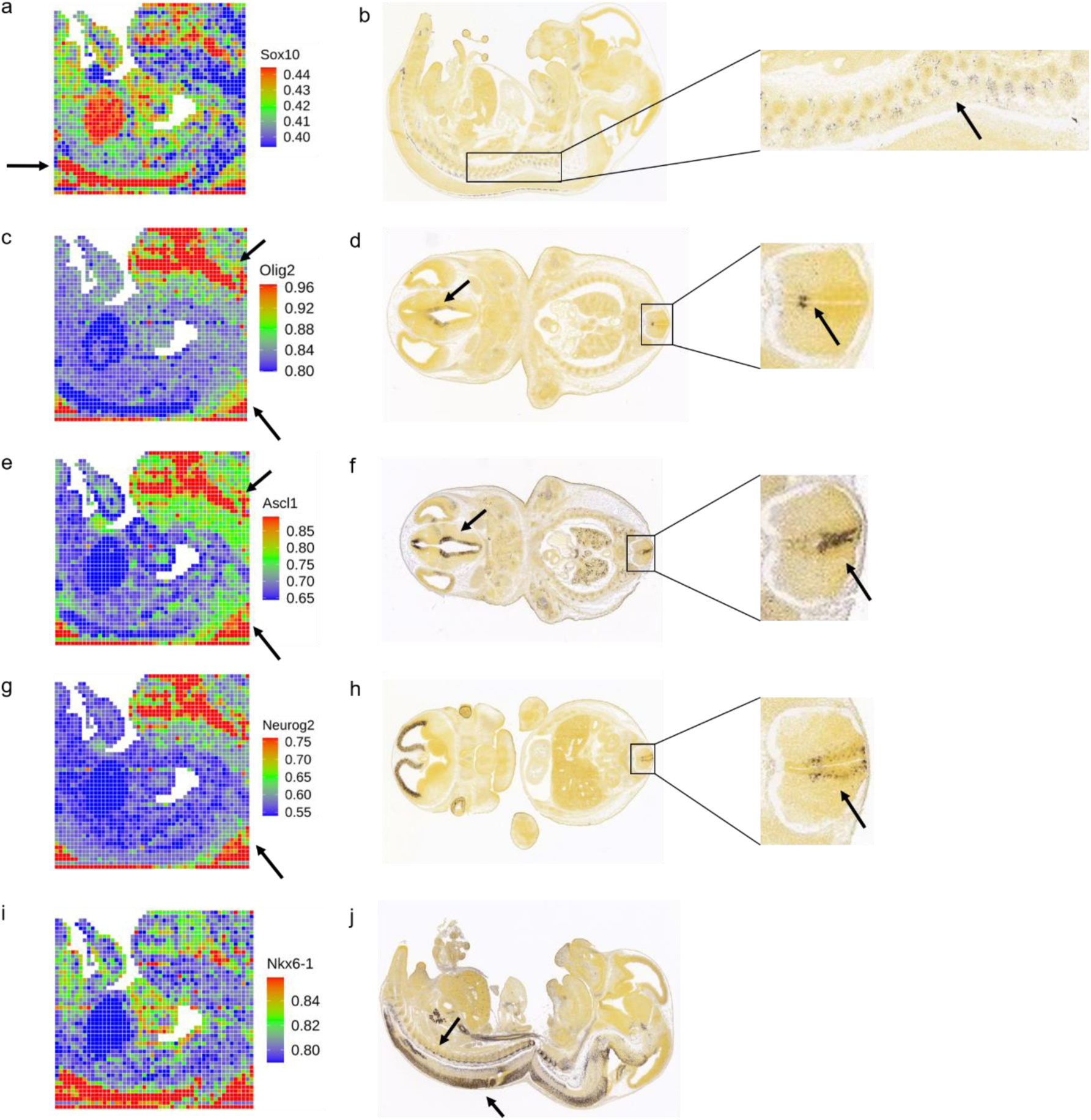
Spatial mapping of gene scores in E13 mouse embryo and comparison with ISH reference data. **a, c, e, g, i,** Spatial mapping of the gene score for selected genes in E13 mouse embryo. **b, d, f, h, j,** In situ hybridization of selected genes at E13.5 mouse embryo from Allen Developing Mouse Brain Atlas.

**Fig. S5.**
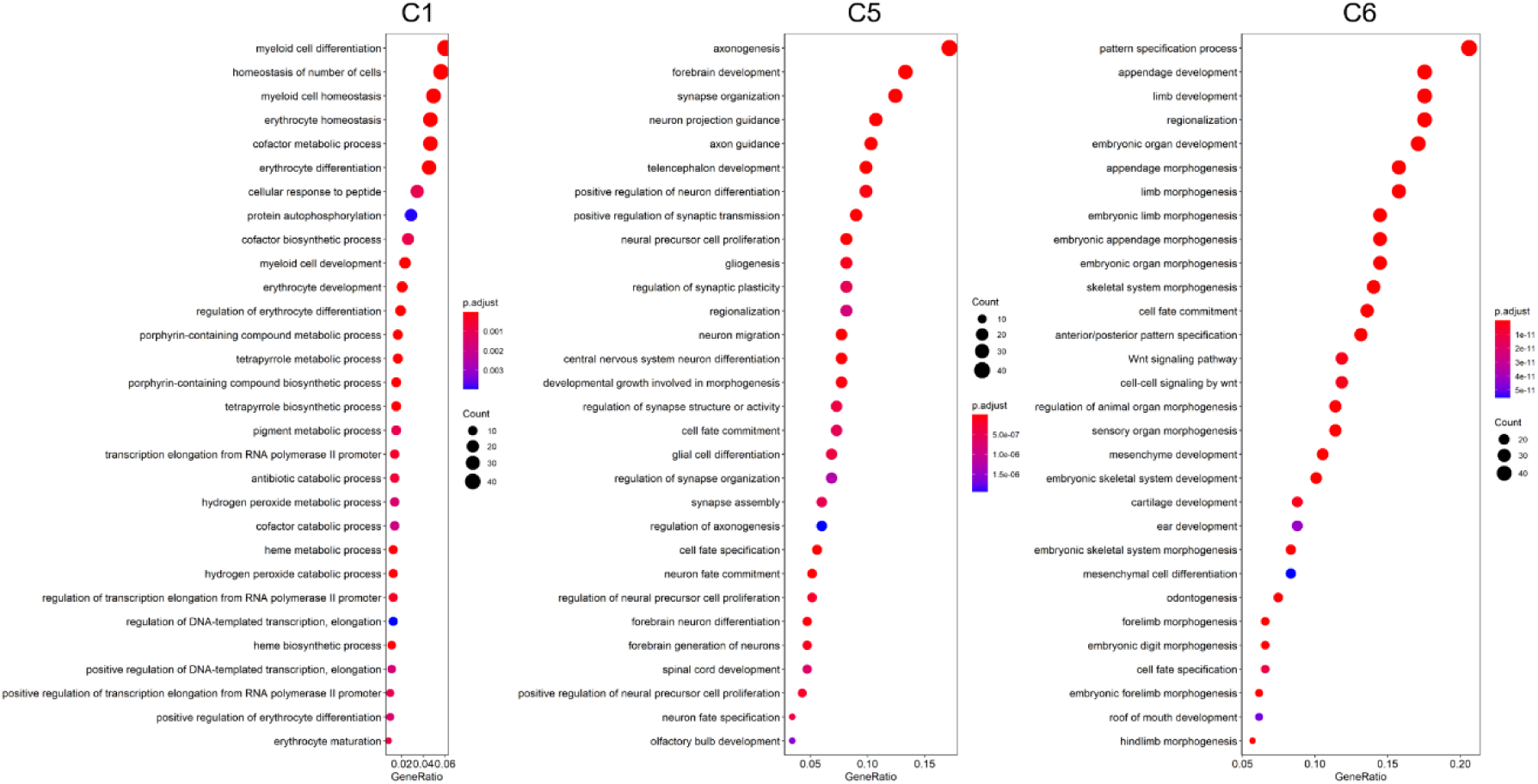
GO enrichment analysis of spatial ATAC-seq data for E13 mouse embryo. GO enrichment analysis of differentially activated genes in selected clusters (C1, C5 and C6).

**Fig. S6.**
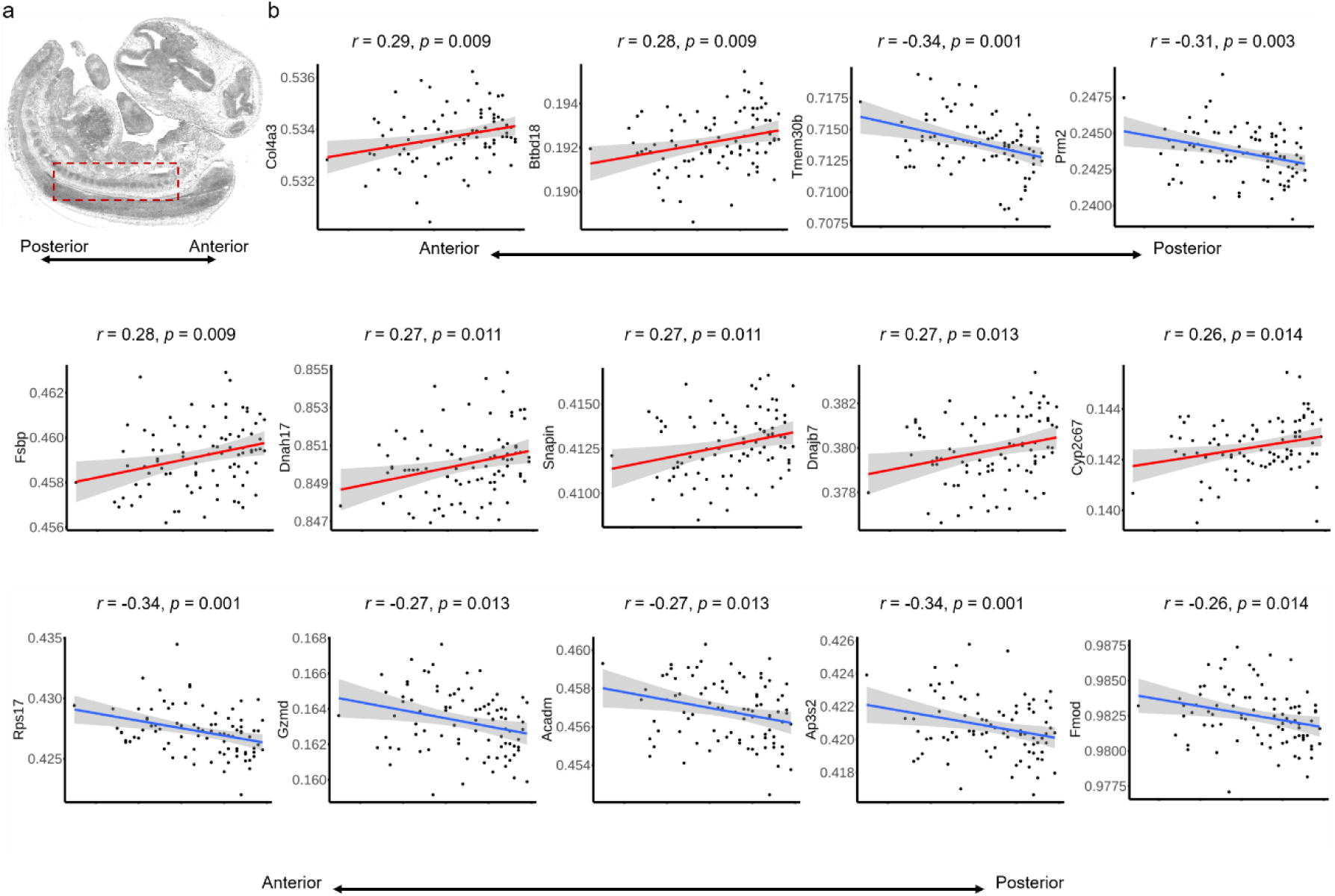
Gene score along the anterior-posterior axis of the spine. **a,** Spine region of E13 mouse embryo profiled by spatial ATAC-seq. **b,** Selected genes found to form expression gradients along the anterior-posterior axis.

**Fig. S7.**
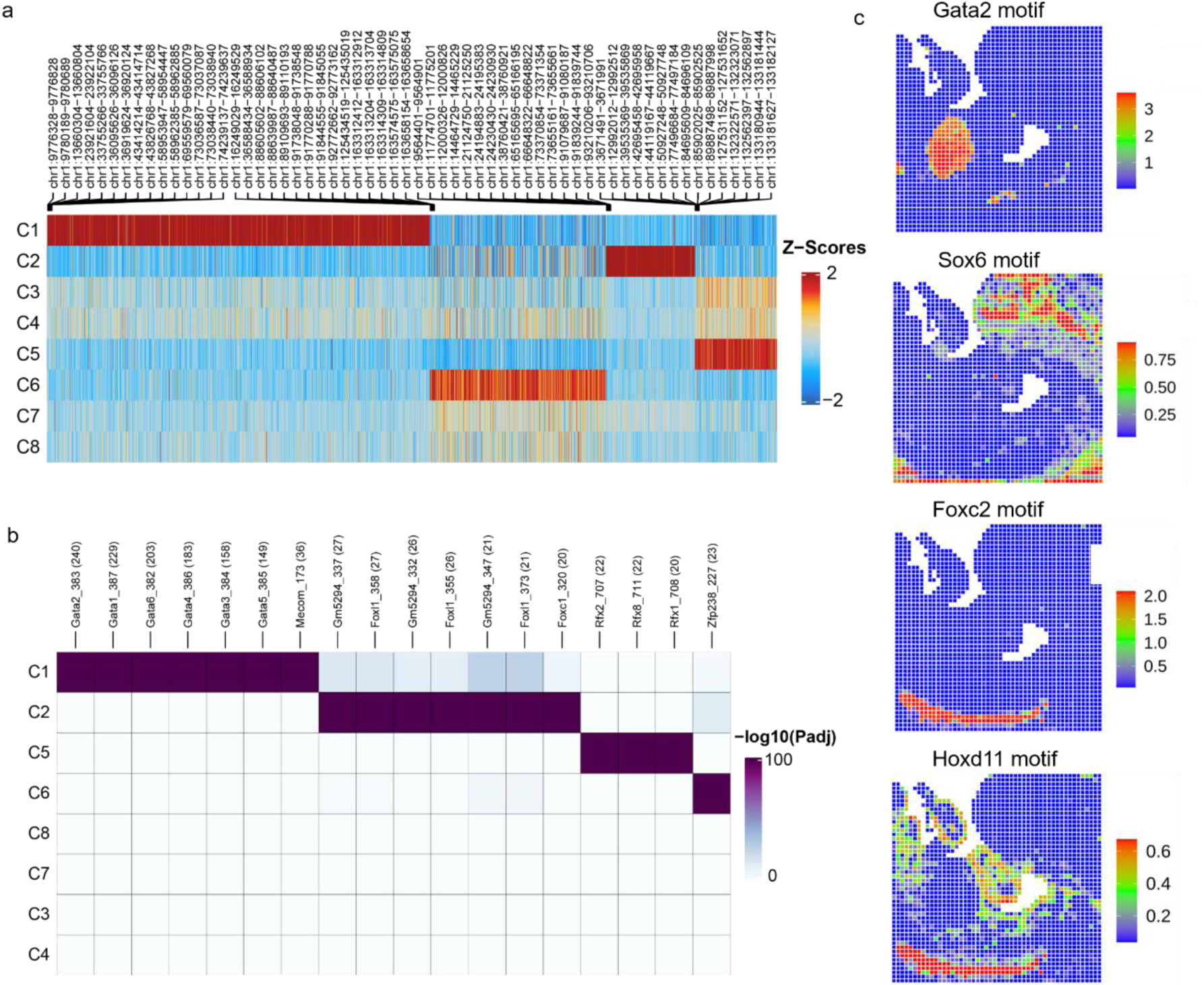
Motif enrichment analysis of the E13 mouse embryo data. **a,** Heatmap of spatial ATAC-seq marker peaks across all clusters identified with bias-matched differential testing. **b,** Heatmap of motif hypergeometric enrichment-adjusted P values within the marker peaks of each cluster. **c,** Spatial mapping of selected TF motif deviation scores.

**Fig. S8.**
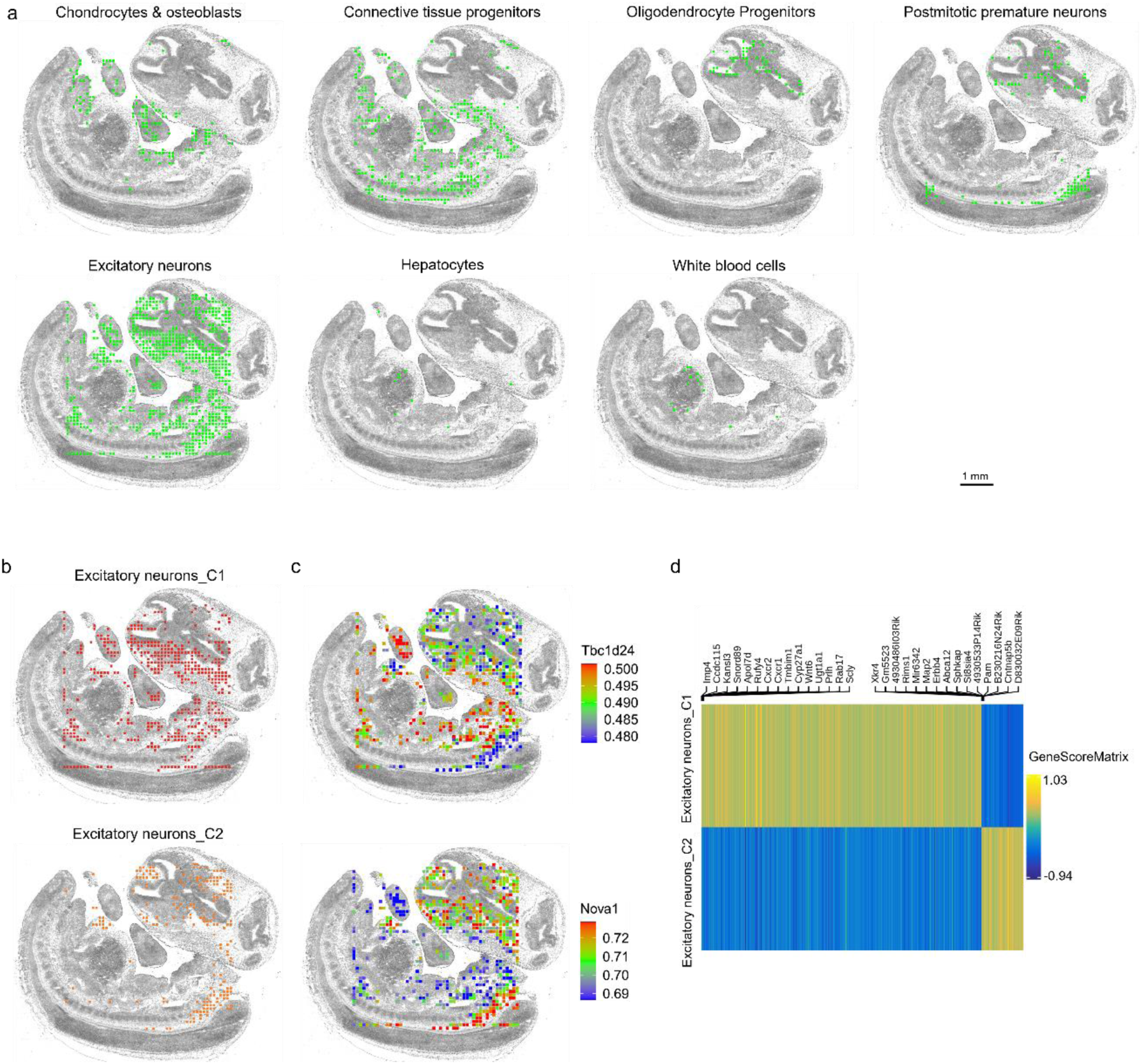
Integrative analysis of spatial ATAC-seq and scRNA-seq for E13 mouse embryo and sub-clustering of excitatory neurons. **a,** Spatial mapping of selected cell types identified by label transferring from scRNA-seq to spatial ATAC-seq. **b-d,** refined clustering process enabled identification of sub-populations in excitatory neurons with distinct spatial distributions (**b**) and marker genes (**c, d**).

**Fig. S9.**
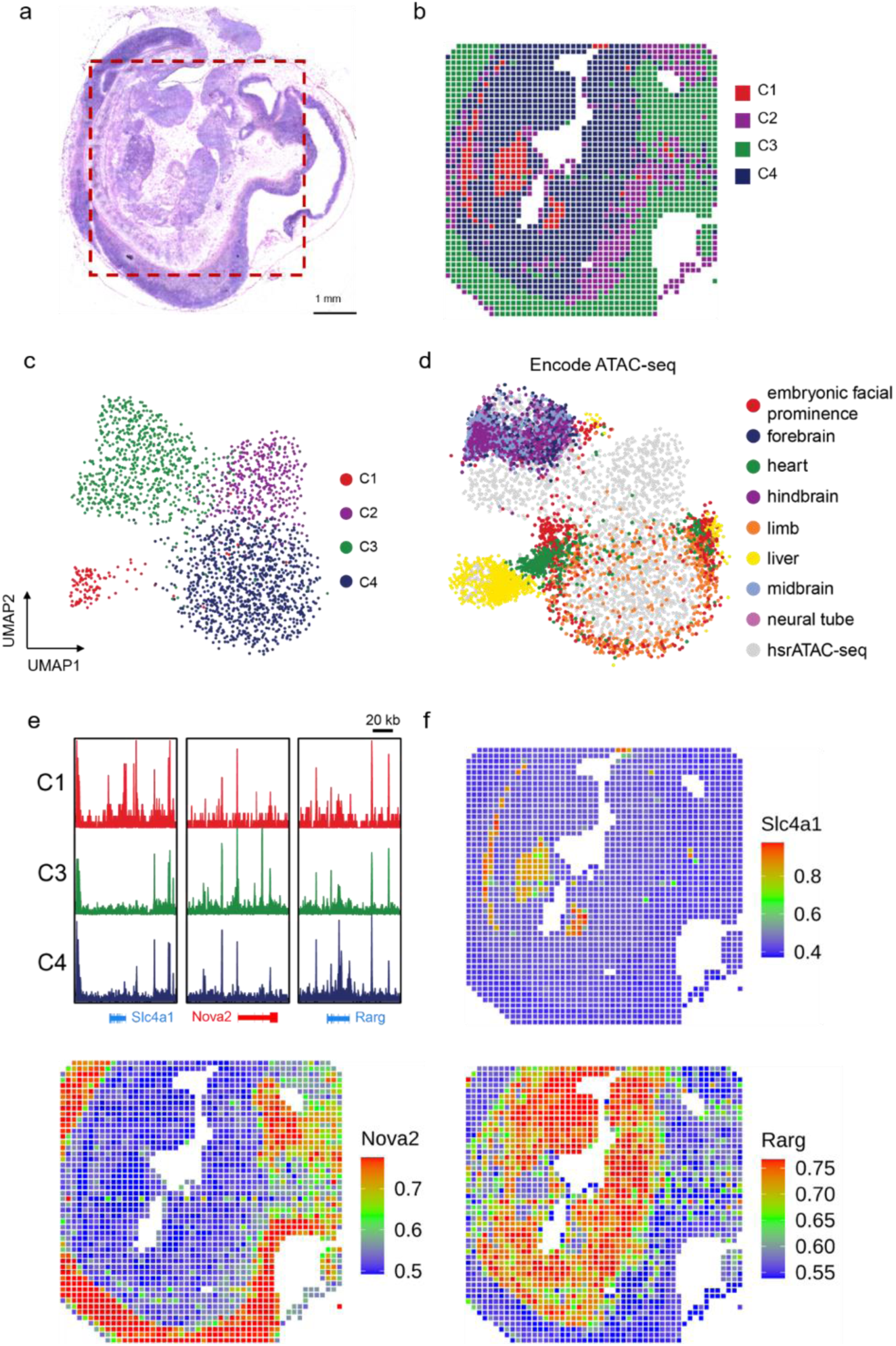
Further analysis of spatial chromatin accessibility mapping of E11 mouse embryo and validation with the ENCODE reference data. **a,** H&E image from an adjacent tissue section and a region of interest for spatial chromatin accessibility mapping (50 µm pixel size). **b,** Unsupervised clustering analysis and spatial distribution of each cluster. **c,** UMAP embedding of unsupervised clustering analysis for spatial ATAC-seq. Cluster identities and coloring of clusters are consistent with (**b**). **d,** LSI projection of ENCODE bulk ATAC-seq data from diverse cell types of the E11.5 mouse embryo dataset onto the spatial ATAC-seq embedding. **e, f,** Genome browser tracks (**e**) and spatial mapping (**f**) of gene scores for selected marker genes in different clusters.

**Fig. S10.**
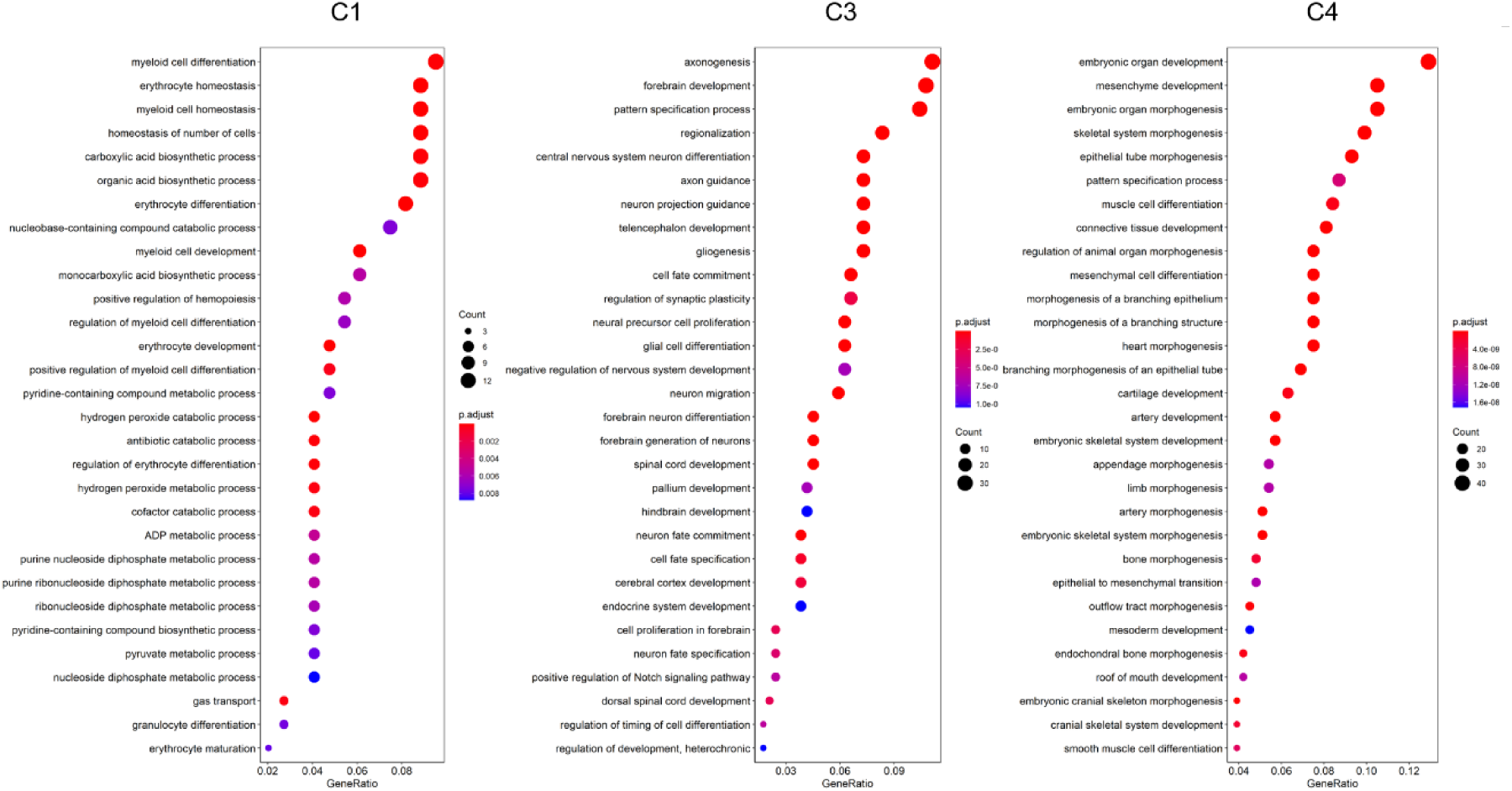
GO enrichment analysis of spatial ATAC-seq data for E11 mouse embryo. GO enrichment analysis of differentially activated genes in selected clusters (C1, C3 and C4).

**Fig. S11.**
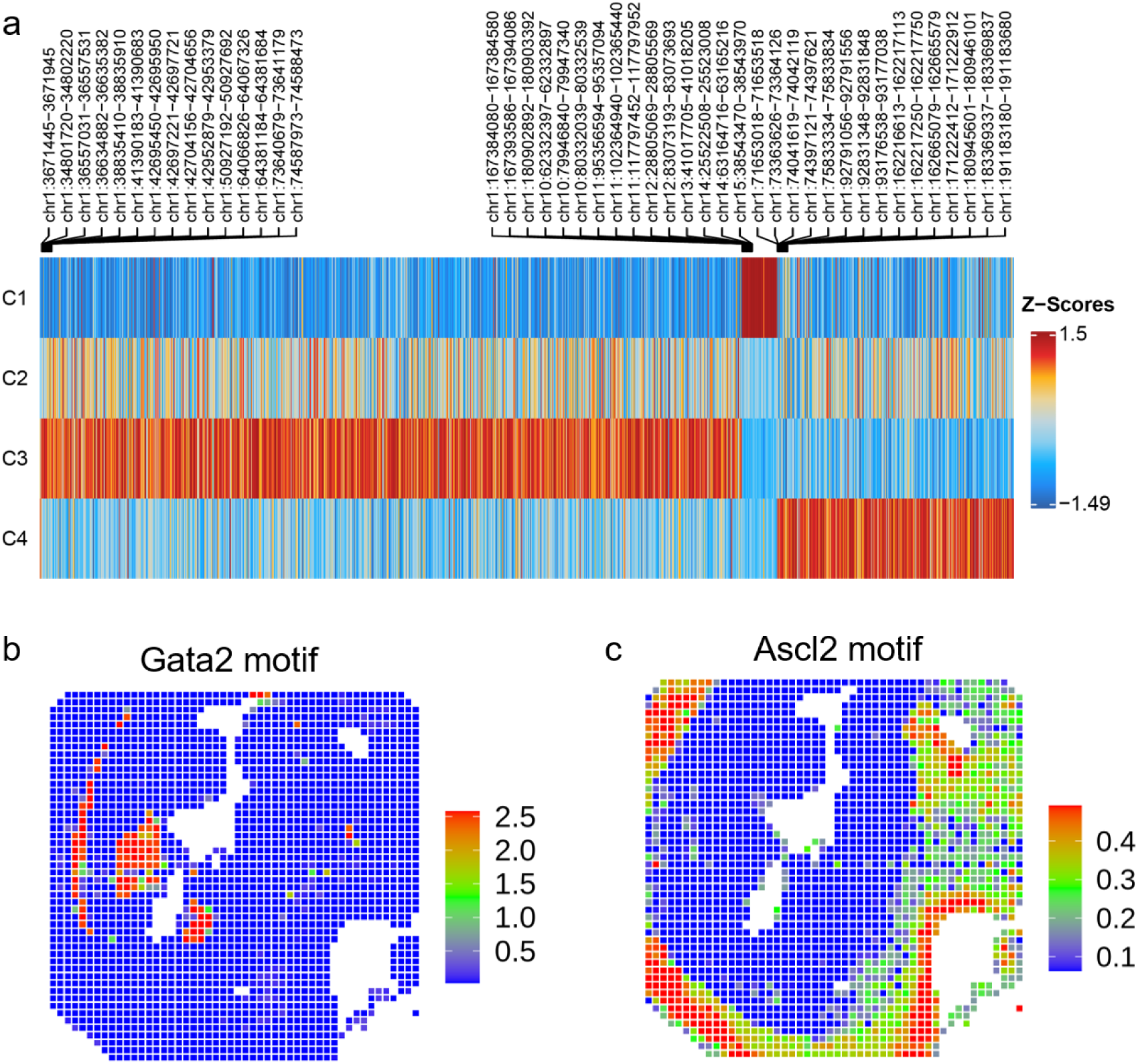
Motif enrichment analysis in E11 mouse embryo. **a,** Heatmap of spatial ATAC-seq marker peaks across all clusters identified with bias-matched differential testing. **b,** Spatial mapping of selected TF motif deviation scores.

**Fig. S12.**
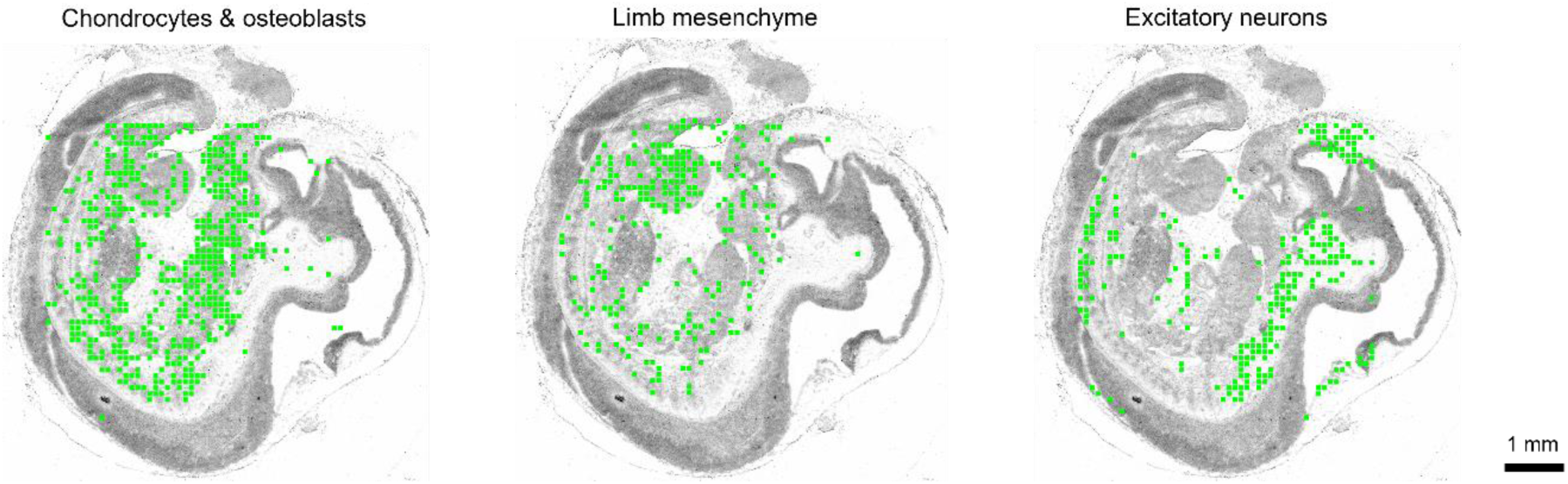
Integrative analysis of spatial ATAC-seq and scRNA-seq for E11 mouse embryo and spatial map visualization of select cell types. Spatial mapping of selected cell types identified by label transferring from scRNA-seq to spatial ATAC-seq.

**Fig. S13.**
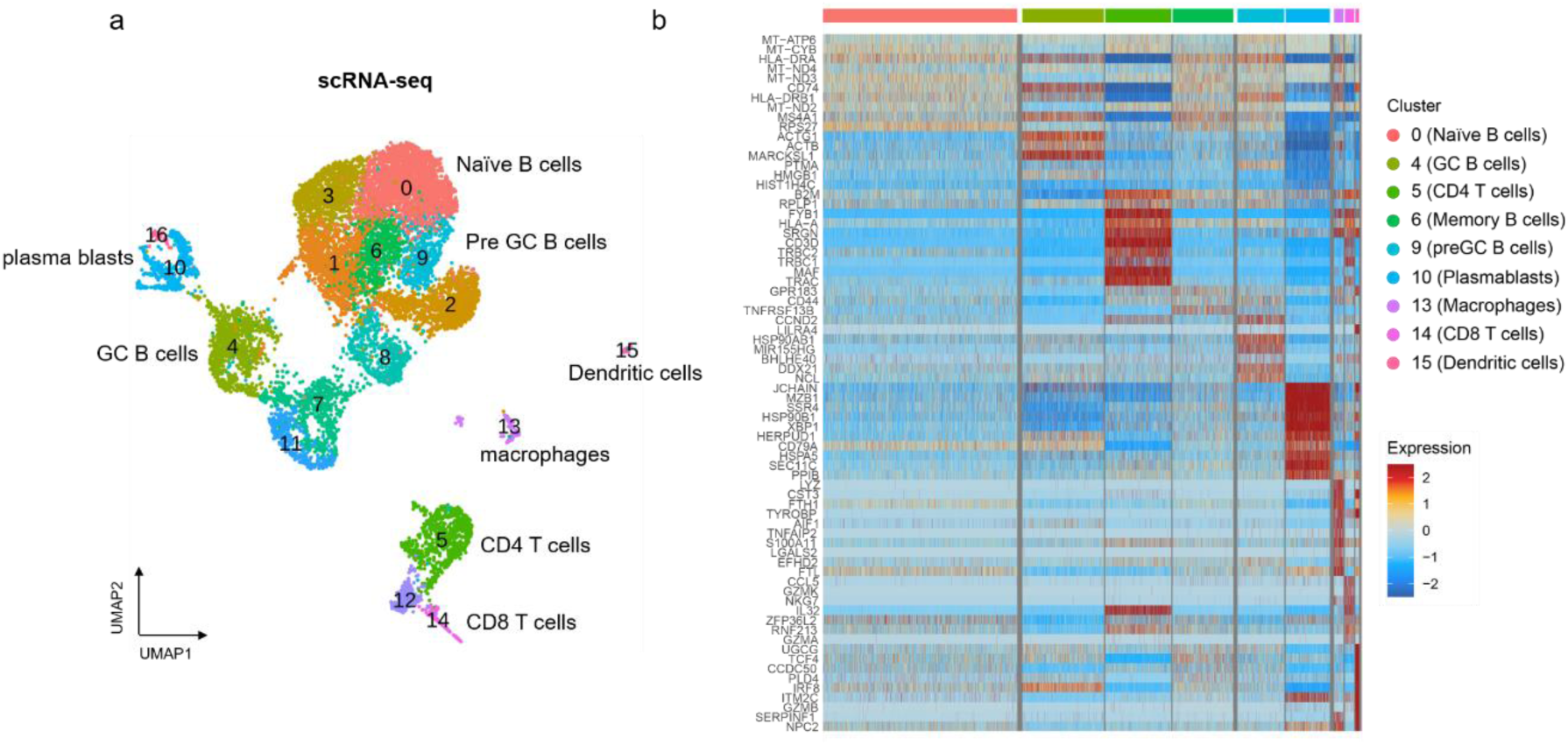
Single-cell mapping of immune cell subsets in human tonsil. **a,** UMAP of tonsillar immune scRNA-seq reference data^33^. **b,** Heatmap comparing key marker gene expression across selected immune cell types.

**Fig. S14.**
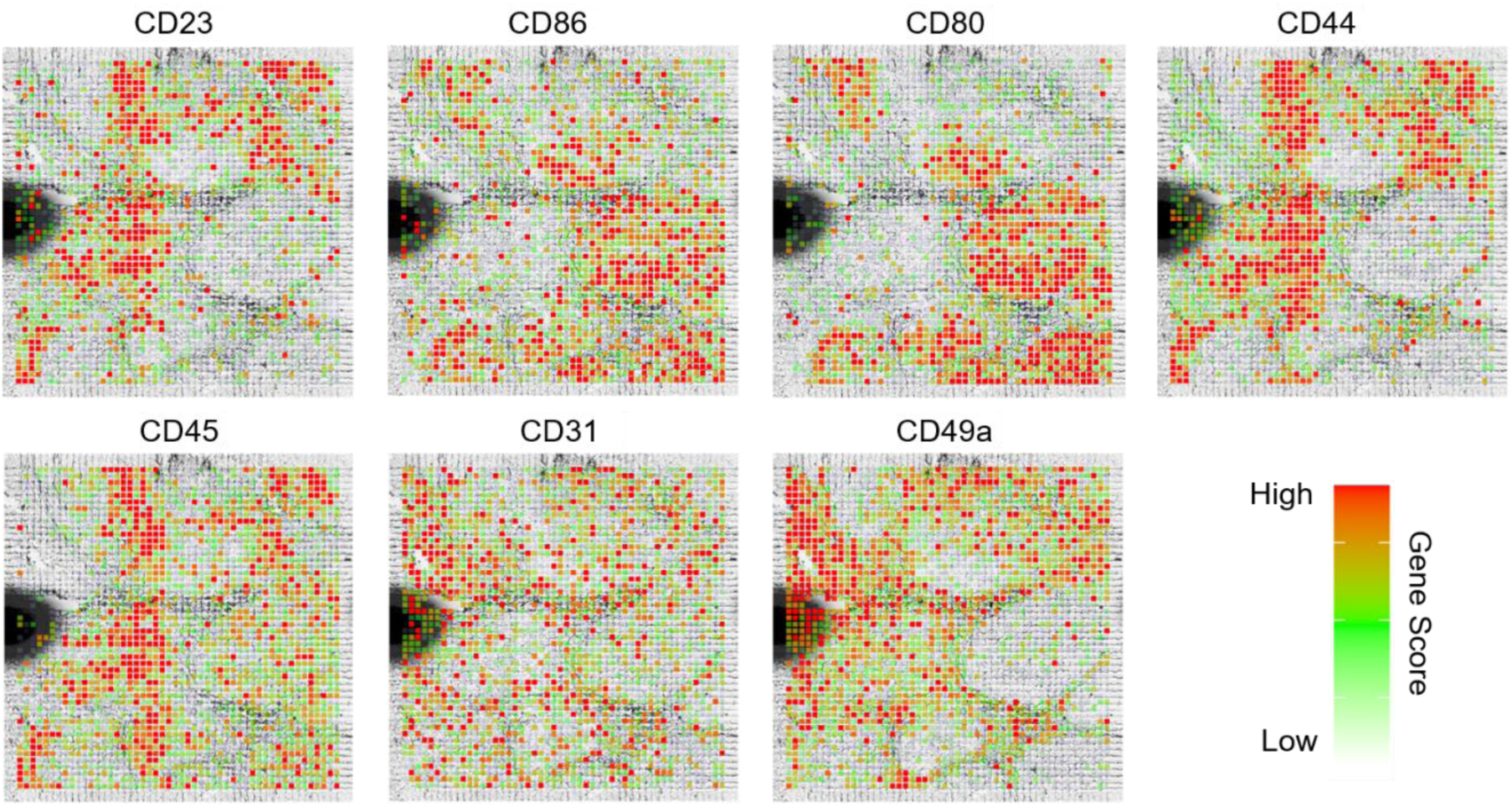
Spatial chromatin accessibility mapping of human tonsil with 20 µm pixel size and visualization of specific marker genes. Spatial mapping of gene scores for selected genes.

**Fig. S15.**
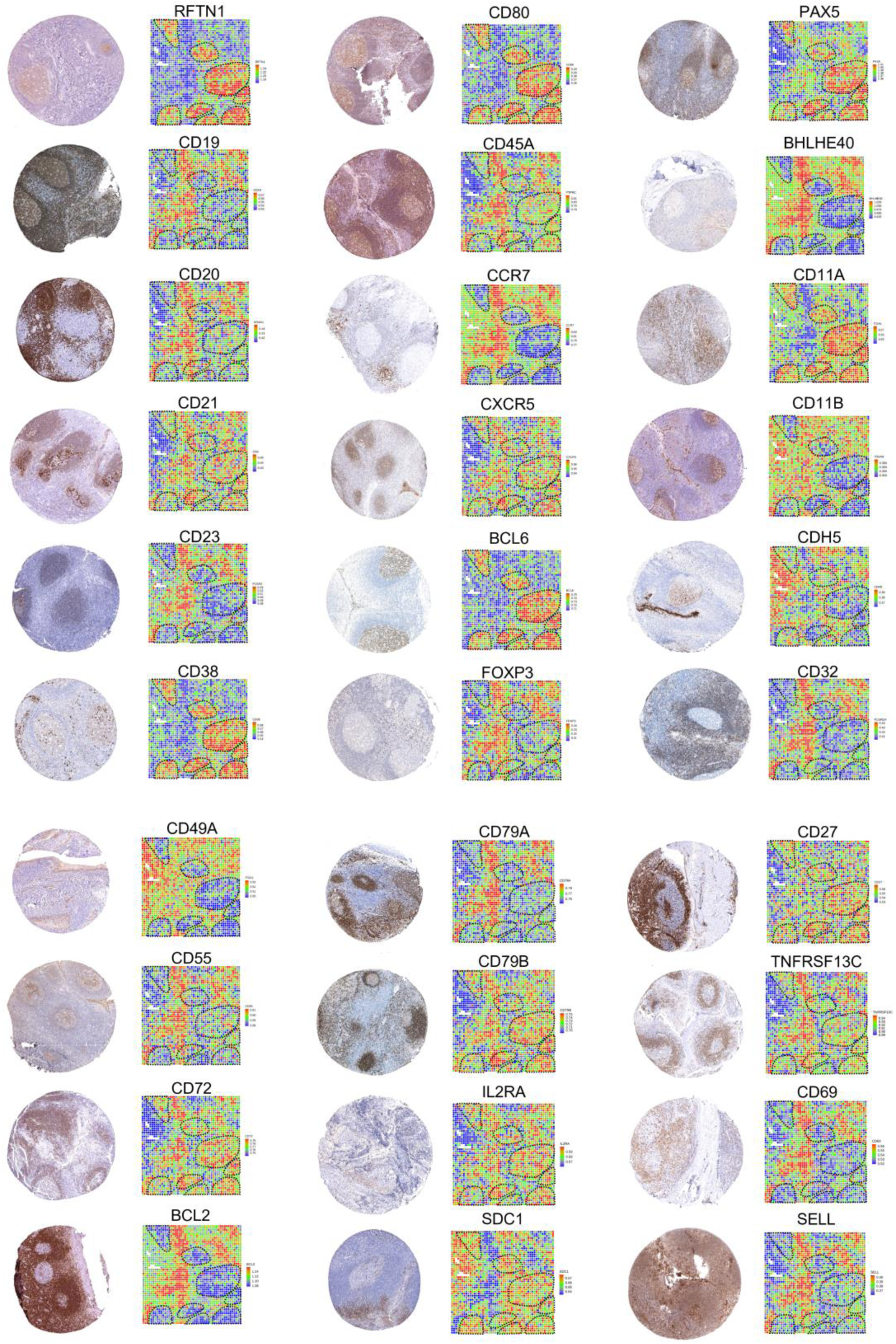
Spatial chromatin accessibility gene score map in comparison with protein expression in human tonsil. The immunohistochemistry reference data were obtained from the Human Protein Atlas^41^.

**Fig. S16.**
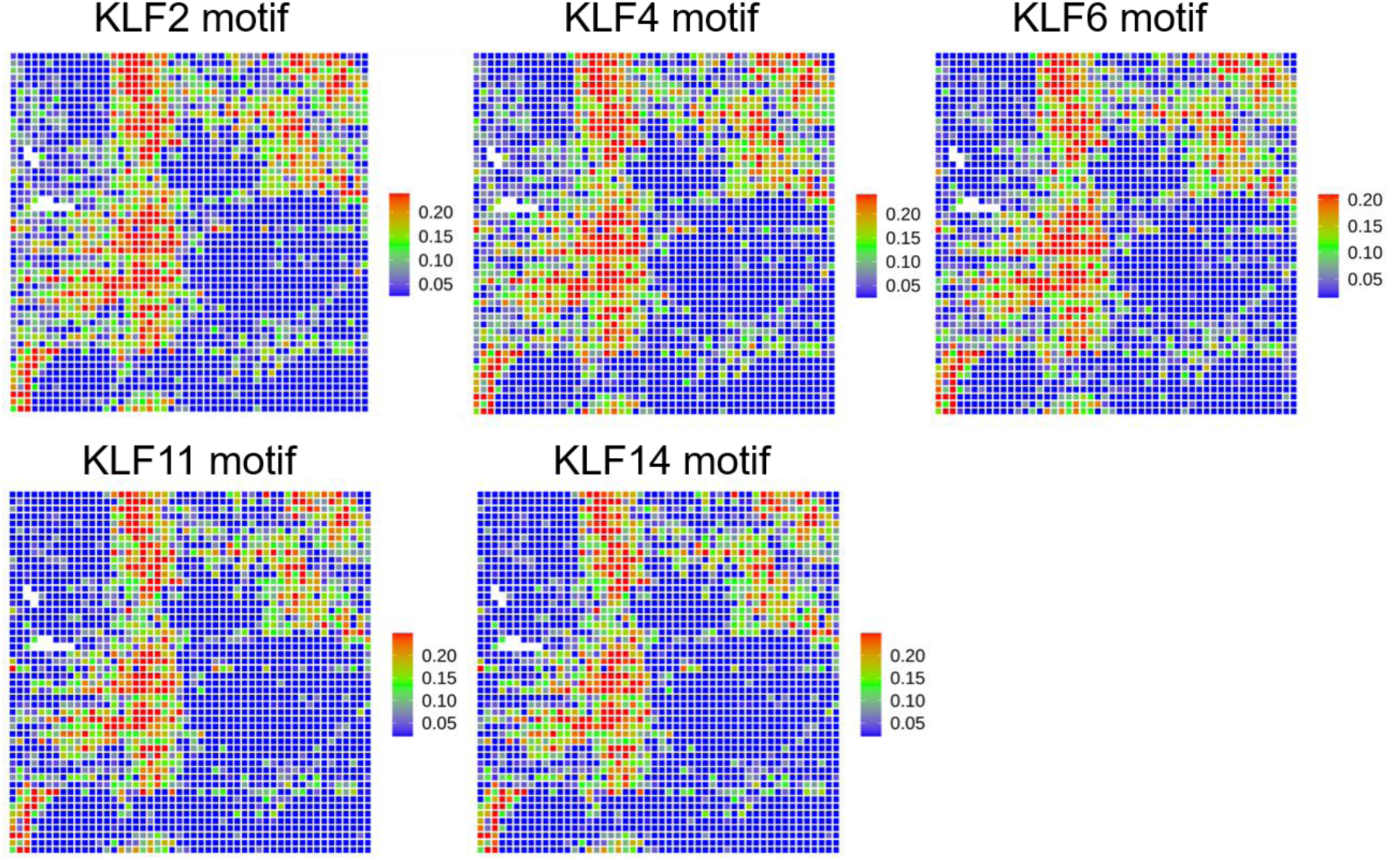
Motif enrichment analysis of spatial ATAC-seq data for human tonsil. Spatial mapping of motif deviation scores for KLF family transcription factors.

**Table S1.**
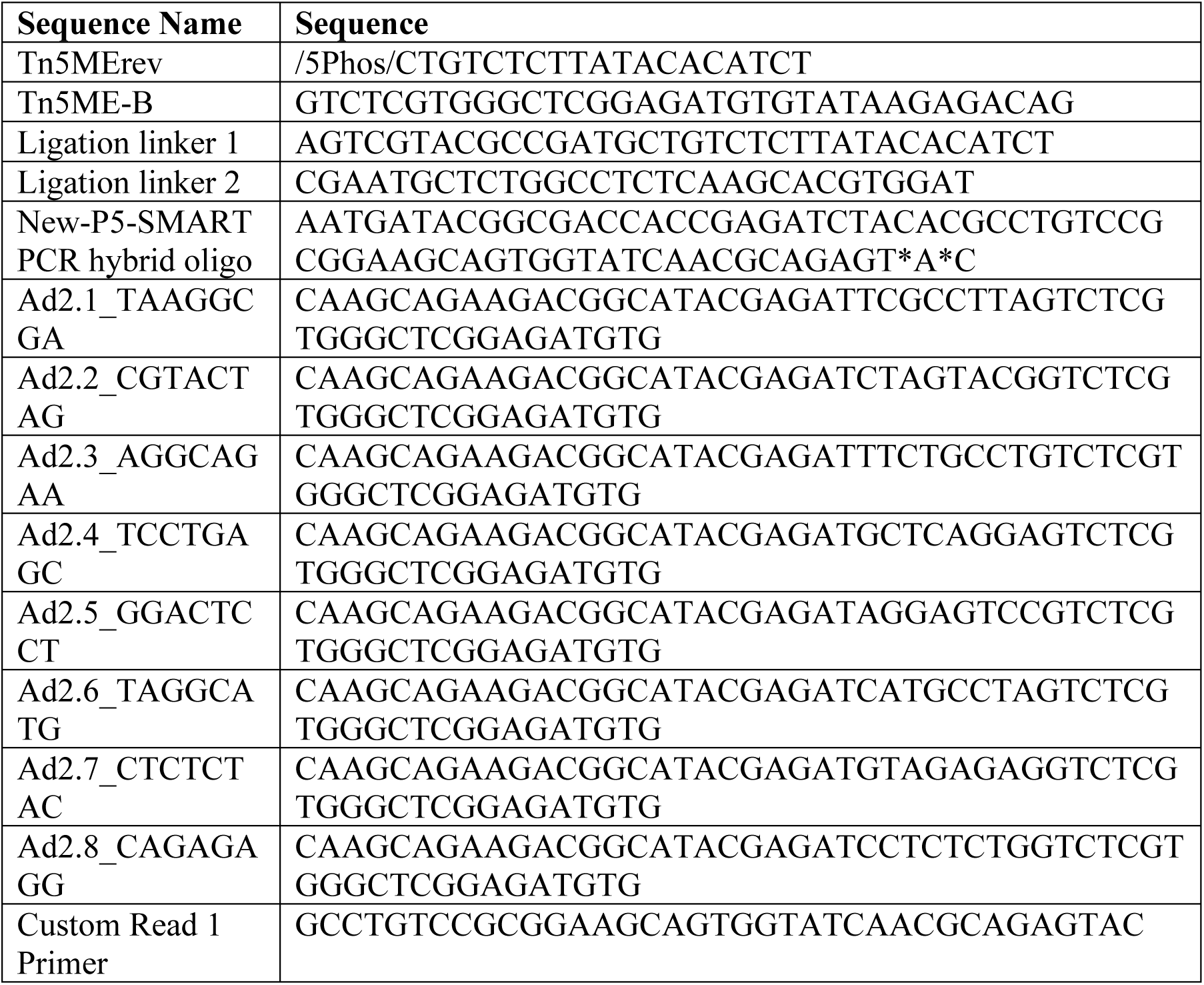
DNA oligos used for PCR and preparation of sequencing library.

**Table S2.**
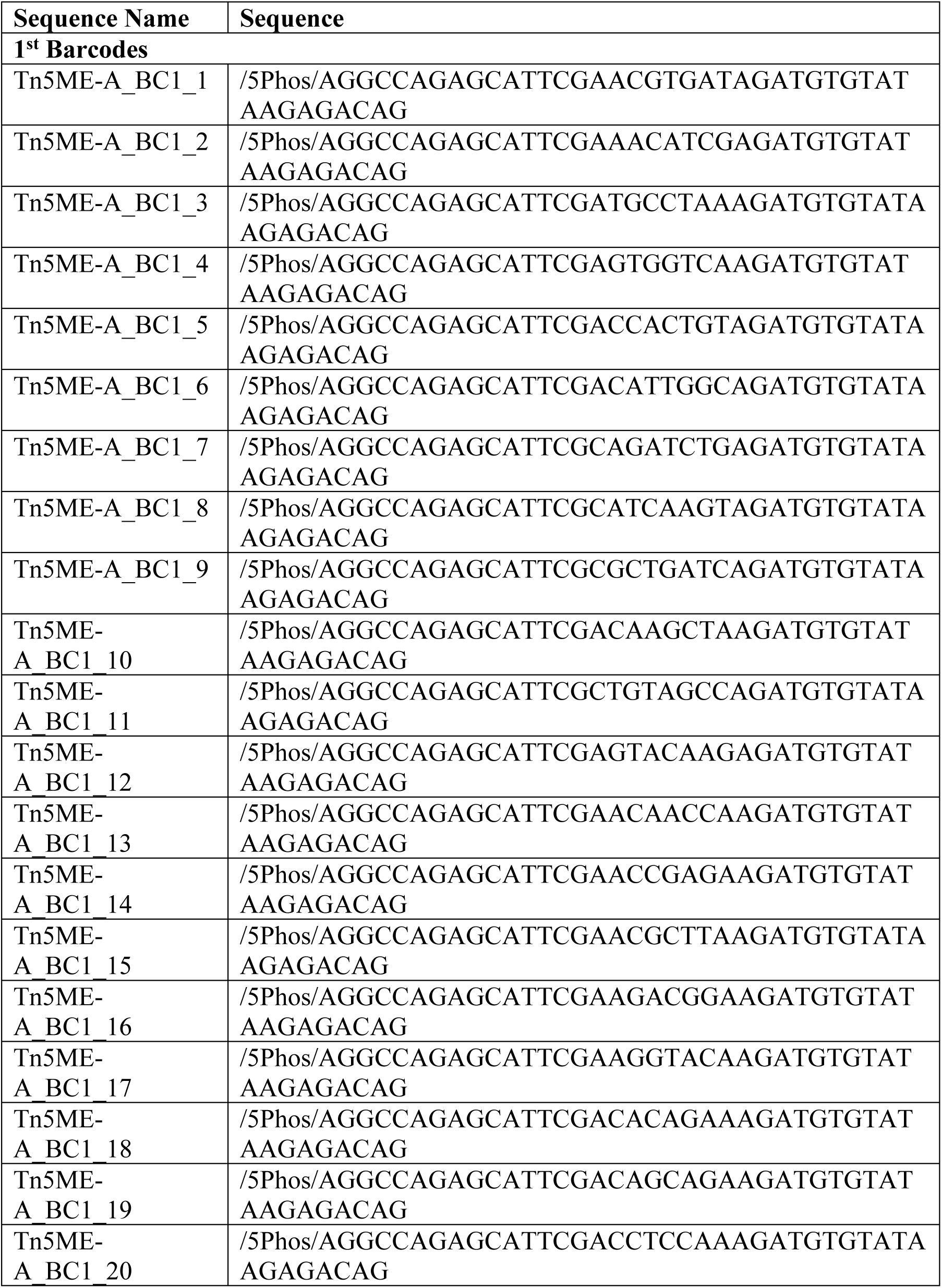
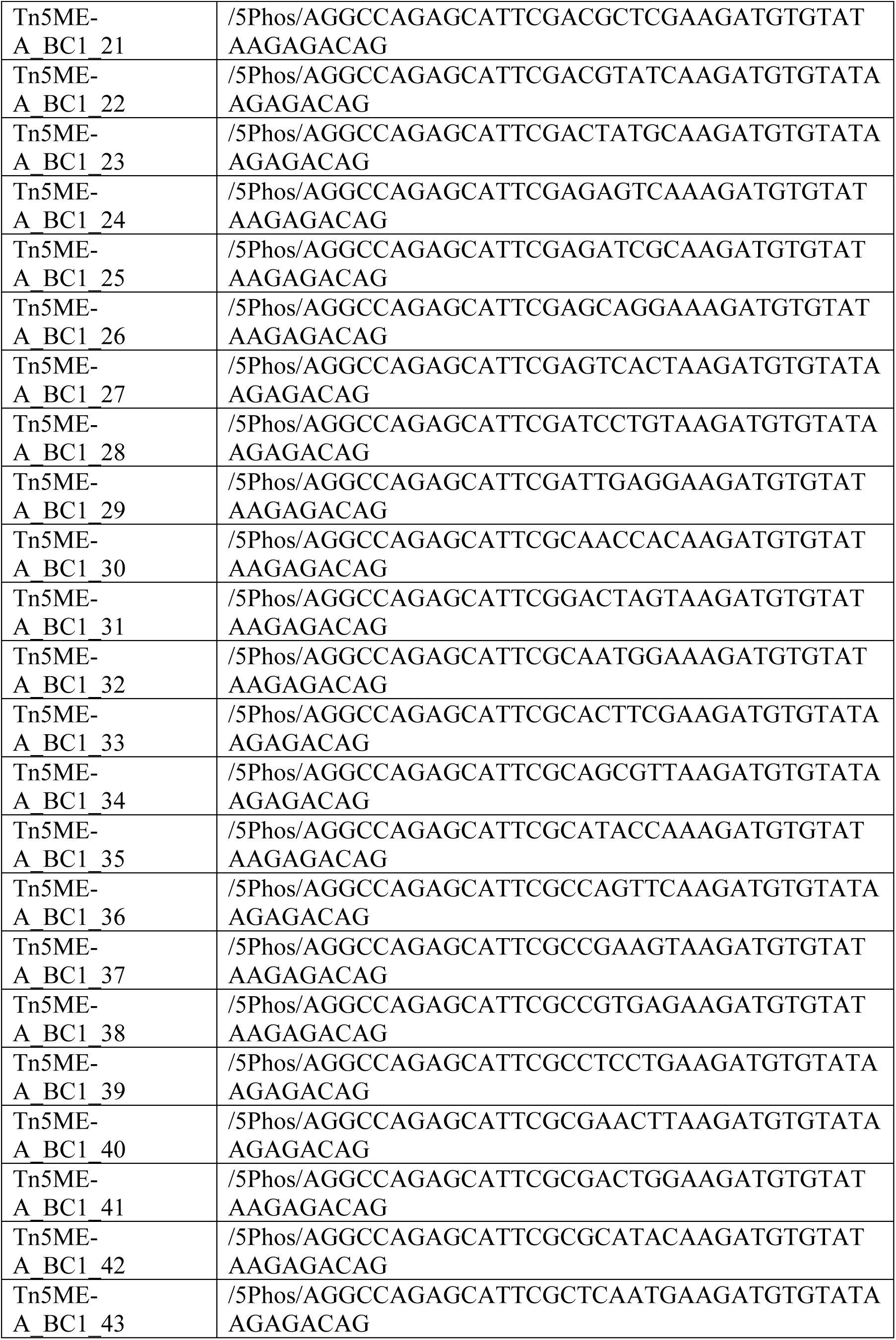
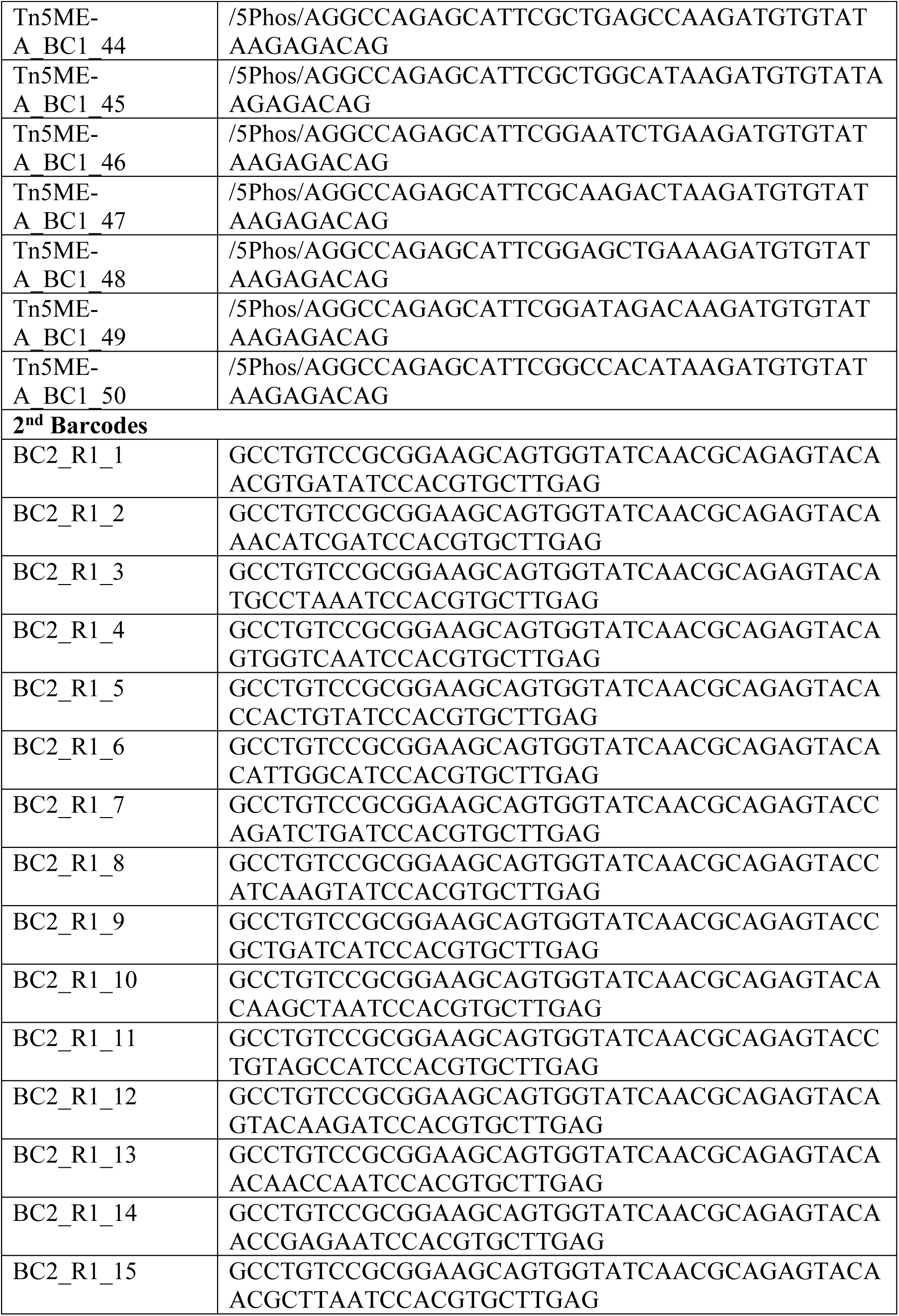
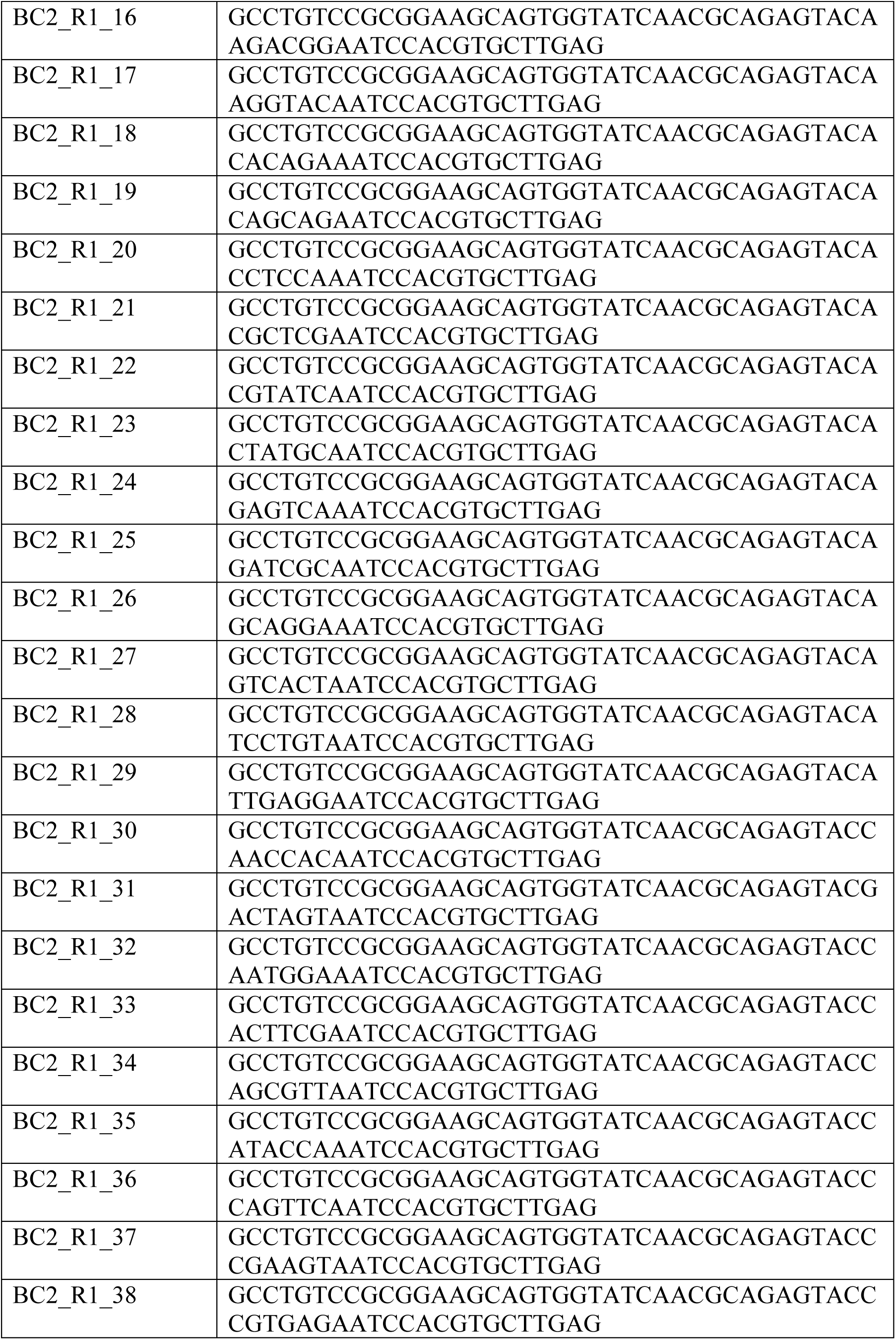
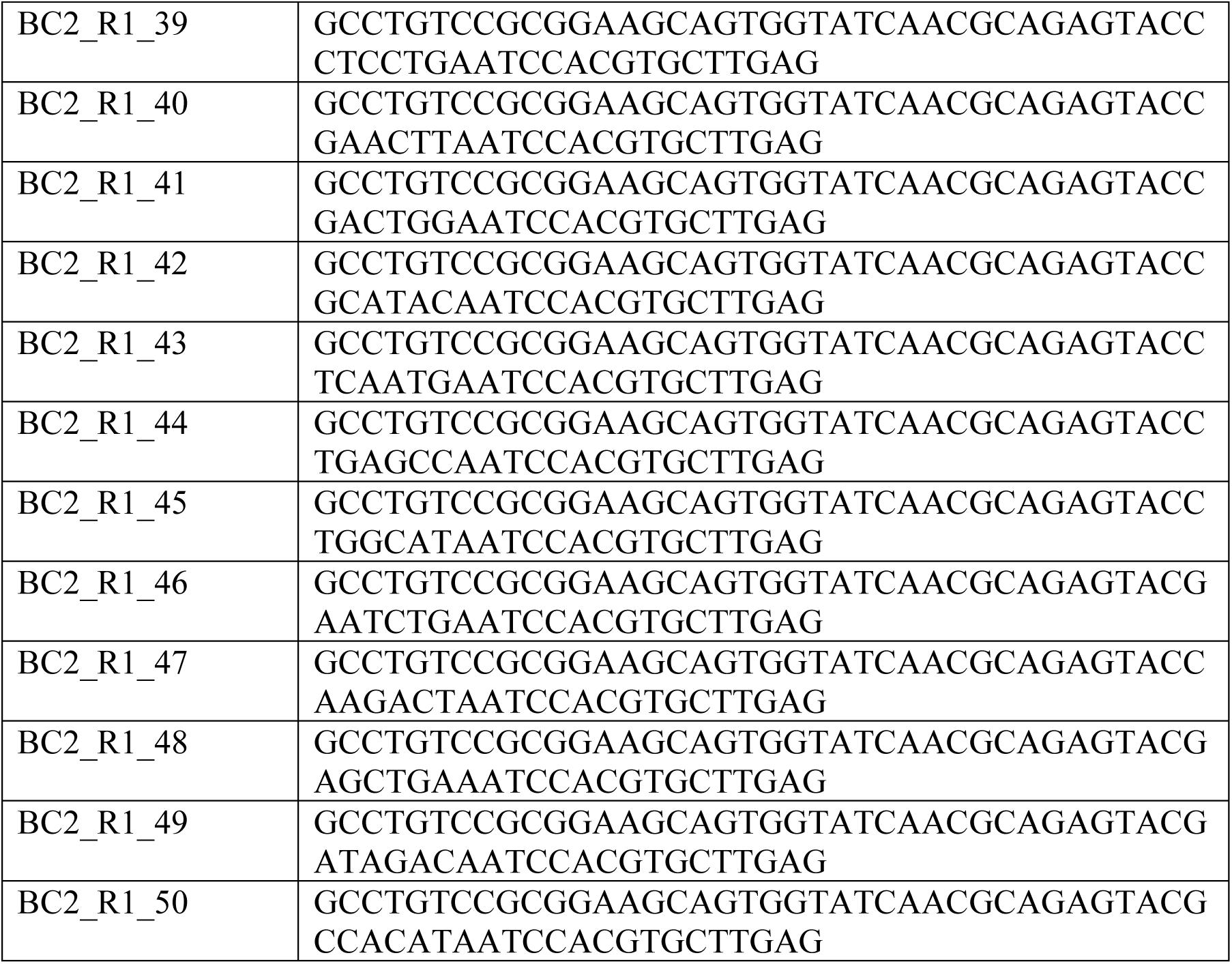
DNA barcode sequences.

**Table S3.**
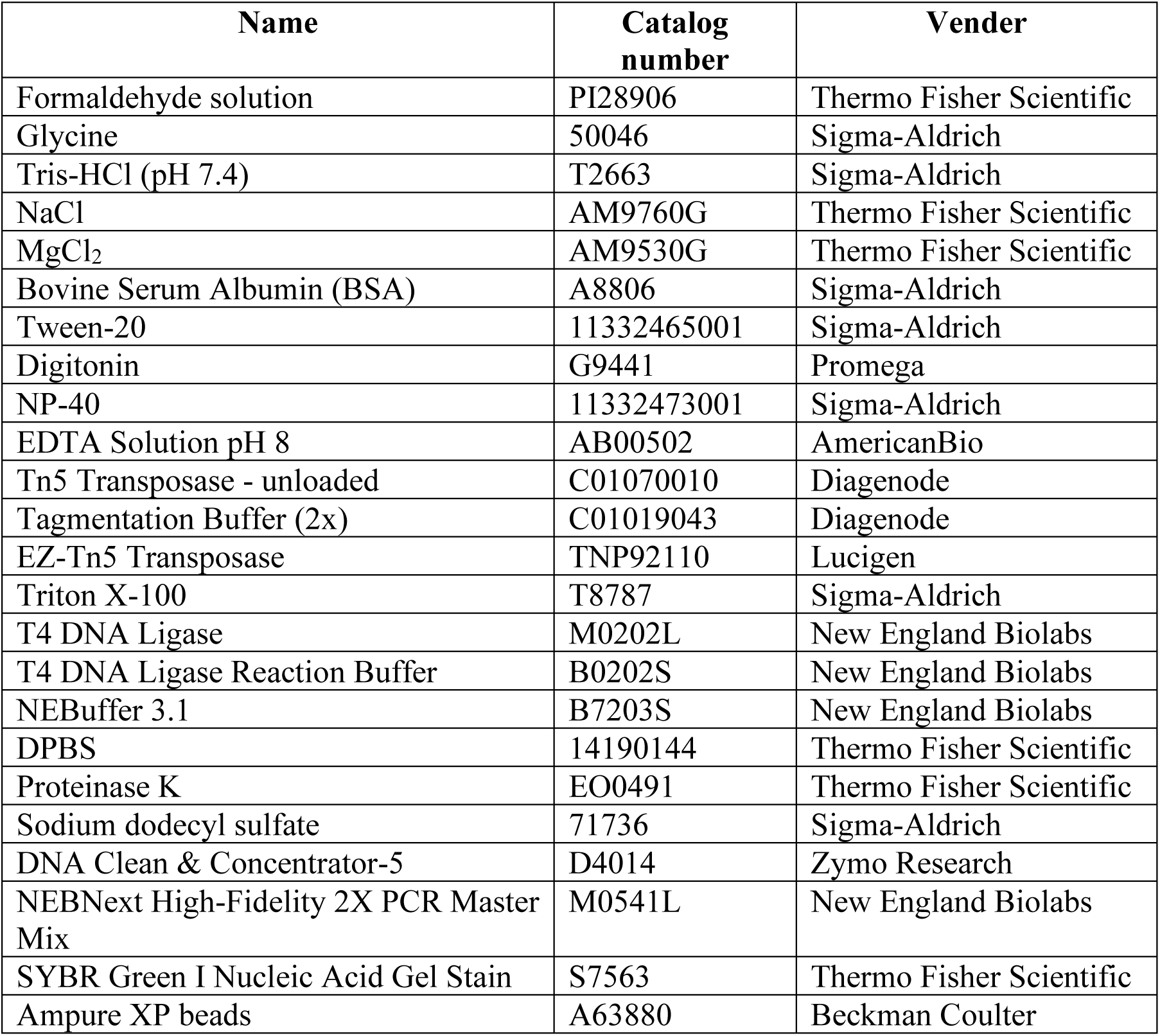
Chemicals and reagents.

